# Phosphoketolase and KDPG aldolase metabolism modulate photosynthetic carbon yield in cyanobacteria

**DOI:** 10.1101/2024.02.12.579767

**Authors:** Ningdong Xie, Chetna Sharma, Katherine Rusche, Xin Wang

## Abstract

Cyanobacteria contribute to roughly a quarter of global net carbon fixation. During diel light/dark growth, dark respiration significantly lowers the overall photosynthetic carbon yield in cyanobacteria and other phototrophs. Currently, it is unclear how respiratory pathways participate in allocating carbon resources at night to optimize dark survival and support daytime photosynthesis. Here we show in the cyanobacterium *Synechococcus elongatus* PCC 7942 that phosphoketolase is orchestrated in an integrative respiratory network in the dark to best allocate carbon resource for amino acid synthesis and prepare for photosynthesis reinitiation upon photoinduction. We further show that the respiratory Entner-Doudoroff (ED) pathway in *S. elongatus* is incomplete, with its key enzyme 2-keto-3-deoxy-6-phosphogluconate (KDPG) aldolase serving to modulate daytime photosynthesis through an alternative oxaloacetate decarboxylation activity. This activity allows for the bypassing of the tricarboxylic acid (TCA) cycle when ATP/NADPH consumption for biosynthesis is excessive and imbalanced relative to their production by light reactions, thereby preventing relative NADPH accumulation and ensuring optimal photosynthetic carbon yield. Optimizing these metabolic processes offers new opportunities to enhance photosynthetic carbon yield in cyanobacteria and other photosynthetic organisms under diel light/dark cycles.

## Introduction

Cyanobacteria account for approximately a quarter of global net primary production (Field et al., 1998; Flombaum et al., 2013). In their natural environment, cyanobacteria and many other photosynthetic organisms experience diel light/dark growth cycles. During the day, photon energy is converted to chemical energy to support growth and synthesize storage polymers. At night, cellular respiration breaks down these storage compounds to support cell maintenance in the dark (Welkie et al., 2018a). Respiration in the dark thus contributes to significant carbon loss in photosynthetic organisms. In plants, respiration alone can lead to as much as 15-30% of carbon loss, significantly lowering the overall photosynthetic carbon yield (Zhu et al., 2008). Optimization of respiratory metabolism thus could minimize carbon loss and enhance the overall photosynthetic carbon yield (Amthor et al., 2019; Hanson et al., 2023). However, studies on respiration and its relations to photosynthesis are relatively sparse. One recent study showed that some of the most prevalent phototrophic microbial lineages in the surface ocean have several fold higher respiration rates at night than the daytime (Munson-McGee et al., 2022).

In cyanobacteria, glycogen is the main energy storage polymer synthesized under light. Glycogen degradation in the dark supplies simple sugars to the oxidative pentose phosphate (OPP) pathway, generating NADPH for energy production through the electron transport chain (Smith, 1983; Lea-Smith et al., 2016). On the other hand, NADPH also serves as cofactors for many detoxification enzymes to clean up reactive oxygen species (ROS) accumulated in the light (Diamond et al., 2017). Although it is well known about this general scheme of cell maintenance in the dark, the details of cellular metabolic state and energy conservation strategy in the dark are far less clear. For instance, since little DNA replication is occurring at night (Watanabe et al., 2012; Ohbayashi et al., 2013), the metabolic destination of C_5_ sugars derived from the OPP pathway is intriguing to understand.

Dark respiration can further impact daytime photosynthesis. When cyanobacteria enter darkness, enzymes for carbon fixation become inactivated (Tamoi et al., 2005), switching cell metabolism to catabolic reactions. In *Synechococcus elongatus* PCC 7942 (hereafter *S. elongatus*), a relatively stable metabolic profile is observed during the first few hours of the night, in contrast to a more drastic metabolic changes in the circadian transcription factor mutant (Δ*rpaA*) (Diamond et al., 2017), suggesting a tight circadian regulation on dark respiration. After dark respiration, several intermediates in the Calvin-Benson-Bassham (CBB) cycle become limited, requiring replenishment for the optimal reinitiation of the CBB cycle reactions upon light exposure (Shinde et al., 2020). Previously, we observed that unlike other CBB cycle intermediates, 3-phosphoglycerate (3PG) accumulates even after darkness and is maintained at a relatively high level during the photosynthesis initiation period (Shinde et al., 2020). This accumulation is not observed in the glycogen synthesis mutant (Δ*glgC*) (Shinde et al., 2020), indicating the origin of 3PG from glycogen degradation in the dark. Through a non-steady-state metabolic flux analysis, a recent study showed that 3PG accumulated in the dark serves as the precursor for carbon regeneration reactions in the CBB cycle during the immediate photoinduction period (Tanaka et al., 2022). Carbon fixation is delayed until the starting substrate ribulose-1,5-bisphosphate (RuBP) is synthesized through carbon regeneration reactions in the CBB cycle. Moreover, 3PG accumulated in the dark also serves as the electron acceptor to quench energy from light reactions upon photoinduction, thus protecting photosystems from potential photodamage.

Dissecting the respiratory network in the dark is central for improving the overall photosynthetic carbon yield in cyanobacteria and other phototrophs. Currently, the metabolic mechanisms through which cyanobacteria allocate carbon and energy resources for efficient cell maintenance in the dark, and the strategies employed to maximize photosynthetic carbon yield during diel light/dark cycles, remain unclear. Here we show that the Entner-Doudoroff (ED) pathway is incomplete in *S. elongatus*, with the key enzyme 2-keto-3-deoxy-6-phosphogluconate (KDPG) aldolase modulating photosynthetic carbon yield through an alternative decarboxylation activity under diel cycles. Furthermore, we show that cyanobacteria form an integrative respiratory network, coupling the OPP pathway, phosphoketolase metabolism, and the Embden-Meyerhof-Parnas (EMP) pathway, to ensure the optimal carbon allocation for cell maintenance and photosynthesis preparation.

## Results

### Glycogen degradation occurs early in the dark to replenish key intermediate pools

Glycogen metabolism is crucial for the fitness of cyanobacteria under diel light/dark growth (Welkie et al., 2018b). We showed previously that *S. elongatus* glycogen synthesis mutant (Δ*glgC*) accumulated significantly less biomass than the wild type under 8h/16h light/dark growth cycles (Shinde et al., 2020). To better understand how growth is affected by different light/dark periods, we grew wild type and the Δ*glgC* mutant in three diel cycles that resemble light/dark lengths in the natural environment, *i.e.*, 10h/14h, 12h/12h, and 14h/10h light/dark cycles. Under all three growth cycles, the Δ*glgC* mutant accumulated less final biomass compared to the wild type (**Fig. 1A**), suggesting glycogen synthesis as an important carbon and energy reservoir to enhance total photosynthetic carbon yield under diel growth.

**Figure 1.**
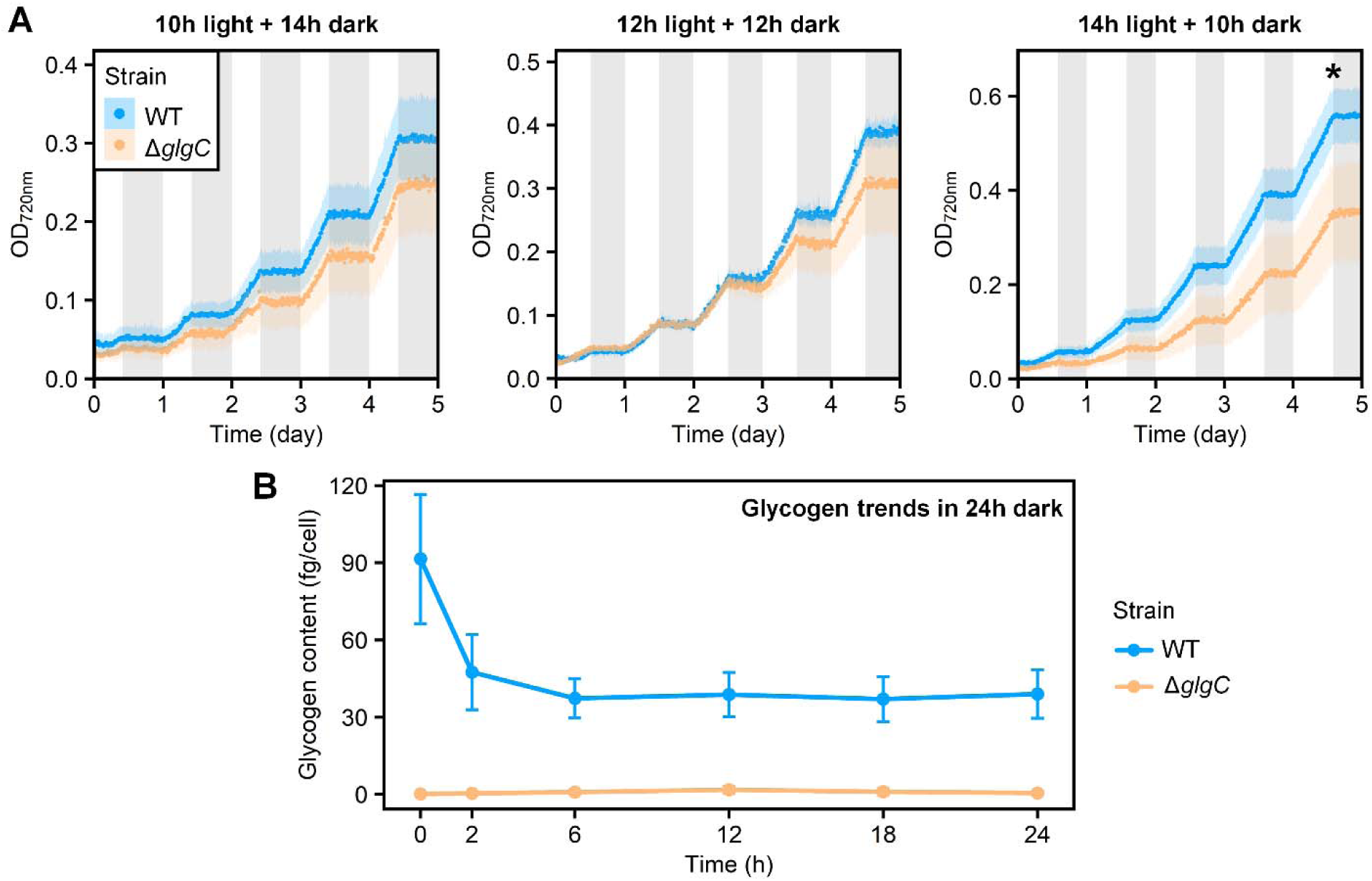
Glycogen degradation primarily occurs early in the dark. **A)** Diel growth of *S. elongatus* wild type (WT) and the glycogen synthesis mutant (Δ*glgC*) during the 10h/14h, 12h/12h, and 14h/10h light/dark cycles (100 µmol/m^2^/s light intensity during the light phase). The dotted line and shading represent the mean and standard deviation, respectively, of the OD_720nm_ measurements for the WT (n = 8) or the Δ*glgC* mutant (n = 4). Grey shading represents the dark period of diel cycles. The terminal OD_720nm_ of WT is higher than that of Δ*glgC* under the 14h/10h light/dark cycle. **B)** Temporal change of the glycogen content in WT and Δ*glgC* during a 24-hour dark period following a continuous growth in the light to late log phase. The circle and error bar represent the mean and standard deviation, respectively, of biological triplicates for both the WT and the Δ*glgC* mutant. Statistical significance was determined using a two-tailed Welch’s t-test. * *p* < 0.05.

To examine how glycogen levels change in the dark, the wild type and Δ*glgC* mutant were shifted to dark after continuous growth in the light to late log phase. No glycogen was detected in the Δ*glgC* mutant, consistent with the inability of this strain to synthesize glycogen (**Fig. 1B**). By contrast, the glycogen synthesized in wild type under light was reduced twofold in abundance during the initial six hours of darkness and remained at constant levels for the duration of the dark period (**Fig. 1B**). Based on our previous study, 3PG is more abundant at the end of the dark period in wild type than Δ*glgC* cells grown in 8h/16h light/dark cycles (Shinde et al., 2020), suggesting that 3PG must be replenished through glycogen degradation during dark respiration. 3PG is a central metabolic intermediate for amino acid synthesis in the dark, and important for carbon fixation reactions upon light exposure (Tanaka et al., 2022). Without glycogen storage, the depletion of 3PG may further disrupt cell metabolism and lead to a higher maintenance cost in the dark.

### Phosphoketolase and KDPG aldolase contribute to optimal photosynthetic carbon yield under diel light/dark growth

In the dark, simple sugars from glycogen degradation are mainly routed to the OPP pathway for energy generation through respiration (Smith, 1983). However, the process by which C_5_ compounds from the OPP pathway will be further catabolized remains unclear. To investigate the potential involvement of other respiratory pathways in the dark, we constructed mutants of the EMP pathway, the phosphoketolase pathway, and the ED pathway by deleting genes for phosphofructokinase (Δ*pfk*), phosphoketolase (Δ*pkt*), and KDPG aldolase (Δ*eda*), respectively (**Supplementary Fig. S1, S2, Table S1**). These mutants were examined for growth under 10h/14h light/dark cycles to maximize the impact of dark respiration. The Δ*pfk* mutant showed similar growth to the wild type, suggesting a negligible role of the upstream EMP pathway in supporting a robust diel growth (**Fig. 2A**). In contrast, both the Δ*pkt* and Δ*eda* single and double mutants exhibited slower growth beginning with the second light phase of the diel cycles and accumulated less terminal biomass compared to the wild type (**Fig. 2B**). By comparison, both the Δ*pfk* and Δ*pkt* mutants grew similarly to the wild type under a full day of continuous light (**Fig. 2, C and D**). The Δ*eda* mutant also showed similar biomass accumulation to the wild type under a 24-h continuous light but had a bleaching phenotype (**Fig. 2, E and F**). These above results suggest that phosphoketolase and KDPG aldolase can modulate carbon budget in cells grown under diel light/dark cycles. Compared to the *S. elongatus* wild type, the Δ*pkt* and Δ*eda* mutants likely have disrupted intermediate pools and lower carbon fixation under diel light/dark growth cycles.

**Figure 2.**
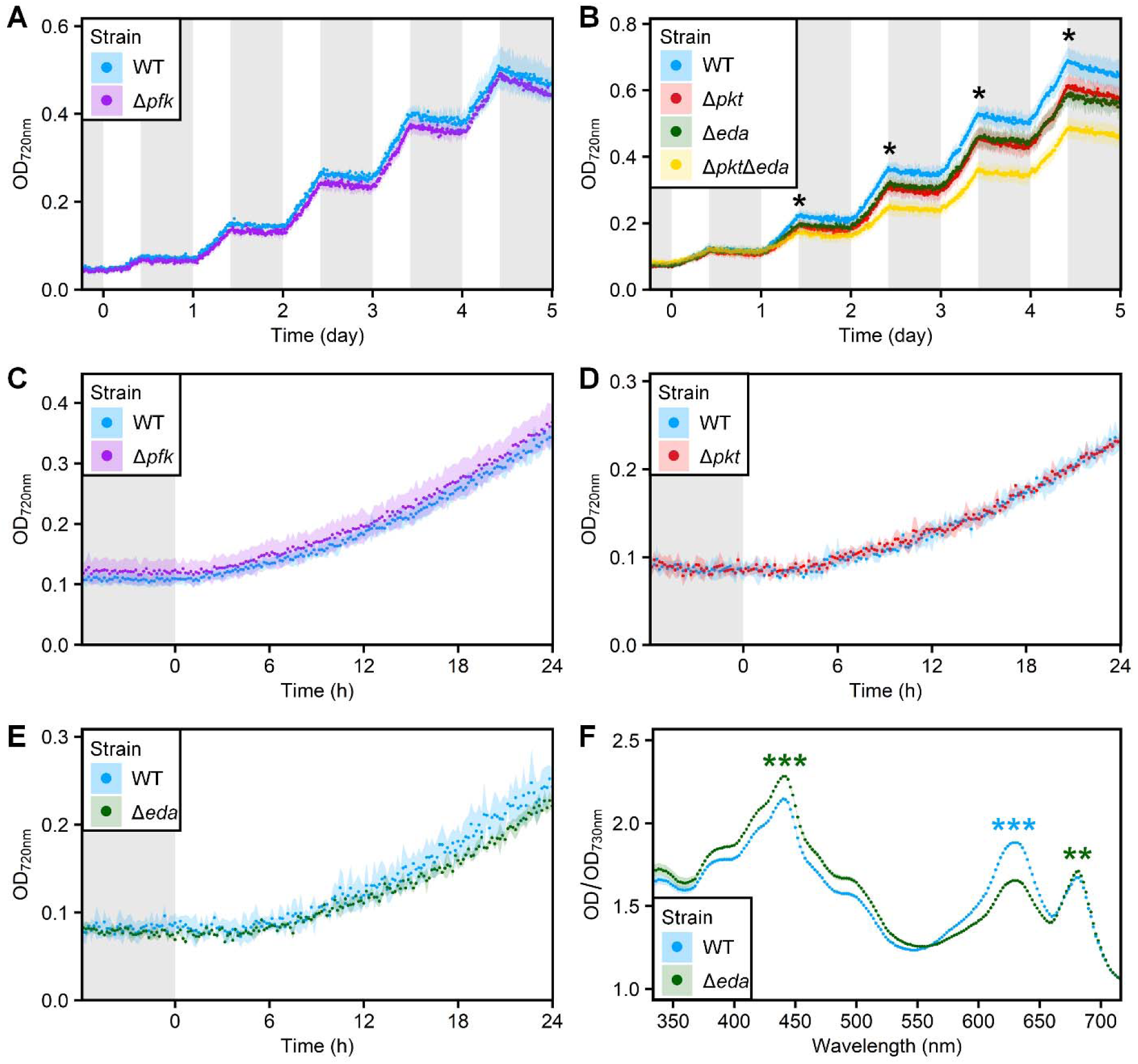
Phosphoketolase and KDPG aldolase contribute to robust diel growth. **A)** *S. elongatus* wild type (WT) and the phosphofructokinase mutant (Δ*pfk*) exhibited similar diel growth during 10h/14h light/dark cycles (*p* = 0.619). **B)** The diel growth of the phosphoketolase mutant (Δ*pkt*), the KDPG aldolase mutant (Δ*eda*), and the *pkt*/*eda* double mutant (Δ*pkt*Δ*eda*) during the 10h/14h light/dark cycles, showing lower biomass than the WT after the second light period (*p* < 0.05, one-tailed Welch’s t-test). **C-E)** Under continuous 24-hour light, the WT showed similar growth to the Δ*pfk* (**C,** *p* = 0.422, t-test on the terminal OD_720nm_), the Δ*pkt* (**D,** *p* = 0.803), and the Δ*eda* (**E,** *p* = 0.517) mutants. The light intensity was maintained at 100 µmol/m^2^/s. The dotted lines and shading represent the OD_720nm_ mean and standard deviation of independently cultivated biological replicates (n = 4 in **A** to **D**; n = 2 for WT and n = 3 for Δ*eda* in **E**). Grey shading indicates the dark periods of diel cycles and the initial 6 hours of dark adaptation to reset the circadian clock. **F)** Full scanning of the visible light spectrum revealed beaching of Δ*eda* under continuous light. The dotted lines and shading represent the change in average OD (normalized by OD_730nm_) and standard deviation of biological replicates (n = 2 for WT and n = 3 for Δ*eda*) across wavelengths from 350 nm to 700 nm. The decreased absorbance at OD_630nm_ indicates phycobilin degradation (bleaching) in the Δ*eda* mutant while the increased OD_440nm_ and OD_680nm_ suggest higher chlorophyll *a* content. Statistical significance was determined using a two-tailed Welch’s t-test unless specified otherwise. * *p* < 0.05, ** *p* < 0.01, *** *p* < 0.001

### Phosphoketolase mutant exhibits high respiration rate in the latter half of the dark period

To examine how phosphoketolase and KDPG aldolase participate in dark respiration, we compared the respiration rates of Δ*pkt* and Δ*eda* to the wild type in the dark period of the diel growth. Following a few 10h/14h light/dark cycles, cells were collected in the subsequent dark period for respiration measurements using a Clark-type oxygen electrode. The respiration rates of all strains increased quickly in the first two hours of darkness, corroborating with the high glycogen degradation activities during this early dark period (**Fig. 3, 1B**). Throughout the dark period, the respiration rates of all strains remained relatively high. Notably, the Δ*pkt* mutant sustained a higher respiration rate than the other two strains, particularly during the latter part of the dark period (**Fig. 3**). This result indicates higher carbon loss in the Δ*pkt* mutant compared to the wild type during dark phase of the diel light/dark cycles.

**Figure 3.**
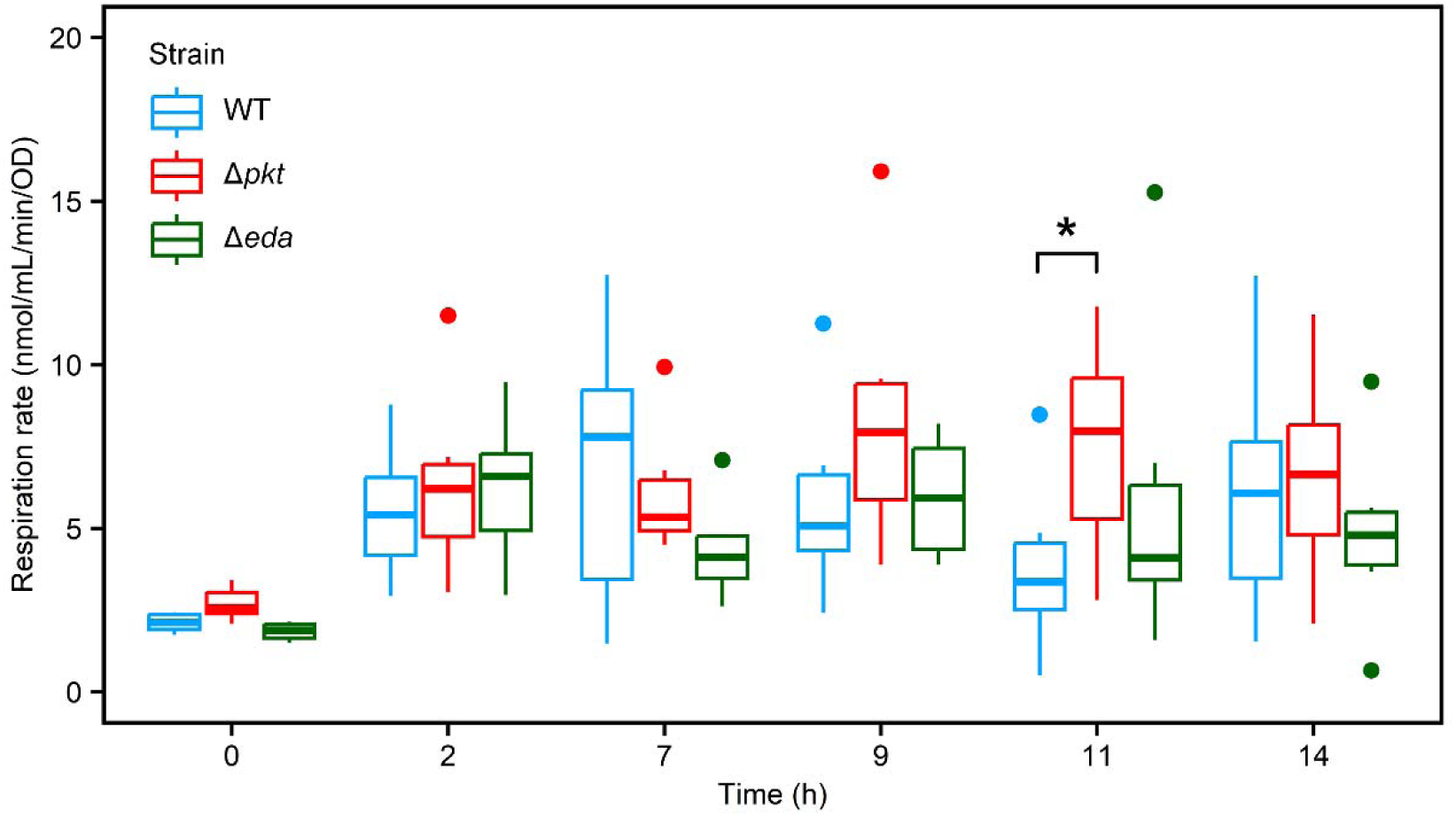
The phosphoketolase mutant exhibit elevated respiration rates in the later dark phase of the diel growth cycles. Respiration rates for the *S. elongatus* wild type (WT), Δ*pkt*, and Δ*eda* mutants were measured with six independently cultivated biological replicates for each strain, collected during the fourth dark period of the 10h/14h light/dark cycle. The data are presented in a box plot format, where the box indicates the 25th and 75th percentiles, the horizontal line within the box denotes the median value, and the whiskers represent the minimum and maximum values, excluding any outliers. Statistical significance was determined using a one-tailed Welch’s t-test. * *p* < 0.05

### KDPG aldolase rather than the ED pathway contributes to robust light/dark growth

In the cyanobacterium *Synechocystis* sp. PCC 6803, the ED pathway was suggested to support cell growth under both mixotrophic conditions in continuous light and autotrophic light/dark growth conditions (Chen et al., 2016). Two key enzymes in the ED pathway are 6-phosphogluconate dehydratase (6PGDH), which converts 6-phosphogluconate into 2-keto-3-deoxy-6-phosphogluconate (KDPG), and KDPG aldolase, which then cleaves KDPG into pyruvate and glyceraldehyde-3-phosphate (G3P). However, no genuine 6PGDH gene has been annotated in the *S. elongatus* genome, and rarely exists in other cyanobacteria (Bachhar and Jablonsky, 2022). Previous research suggests that dihydroxy acid dehydratase (DHAD) (*ilvD*) in amino acid metabolism could serve as an alternative enzyme for the dehydratase activity in the ED pathway in cyanobacteria (Chen et al., 2016). To examine this possibility, we purified the *S. elongatus* DHAD enzyme (*ilvD*) and tested its activity toward the ED pathway intermediate 6PG. Although *S. elongatus* DHAD shares a similar predicted structure to the genuine 6PGDH of other bacteria (**Supplementary Fig. S3**), it showed no 6PG dehydratase activity and instead displayed strong activity toward its native substrate 2,3-dihydroxyisovalerate (**Fig. 4, A and B; Supplementary Fig. S4**). These results suggest that DHAD is not an alternative enzyme for the ED pathway in *S. elongatus*. To examine whether an alternative enzyme could serve as 6PG dehydratase in *S. elongatus*, we conducted *in vitro* 6PGDH enzyme assays by supplementing 6PG to cell lysates of both light and dark incubated cells. However, KDPG, the product of 6PG dehydration, was not detected in these cell lysates, suggesting it is either absent or present at concentrations below the detection limit (**Fig. 4C**). These results strongly suggest that the ED pathway is incomplete in *S. elongatus*.

**Figure 4.**
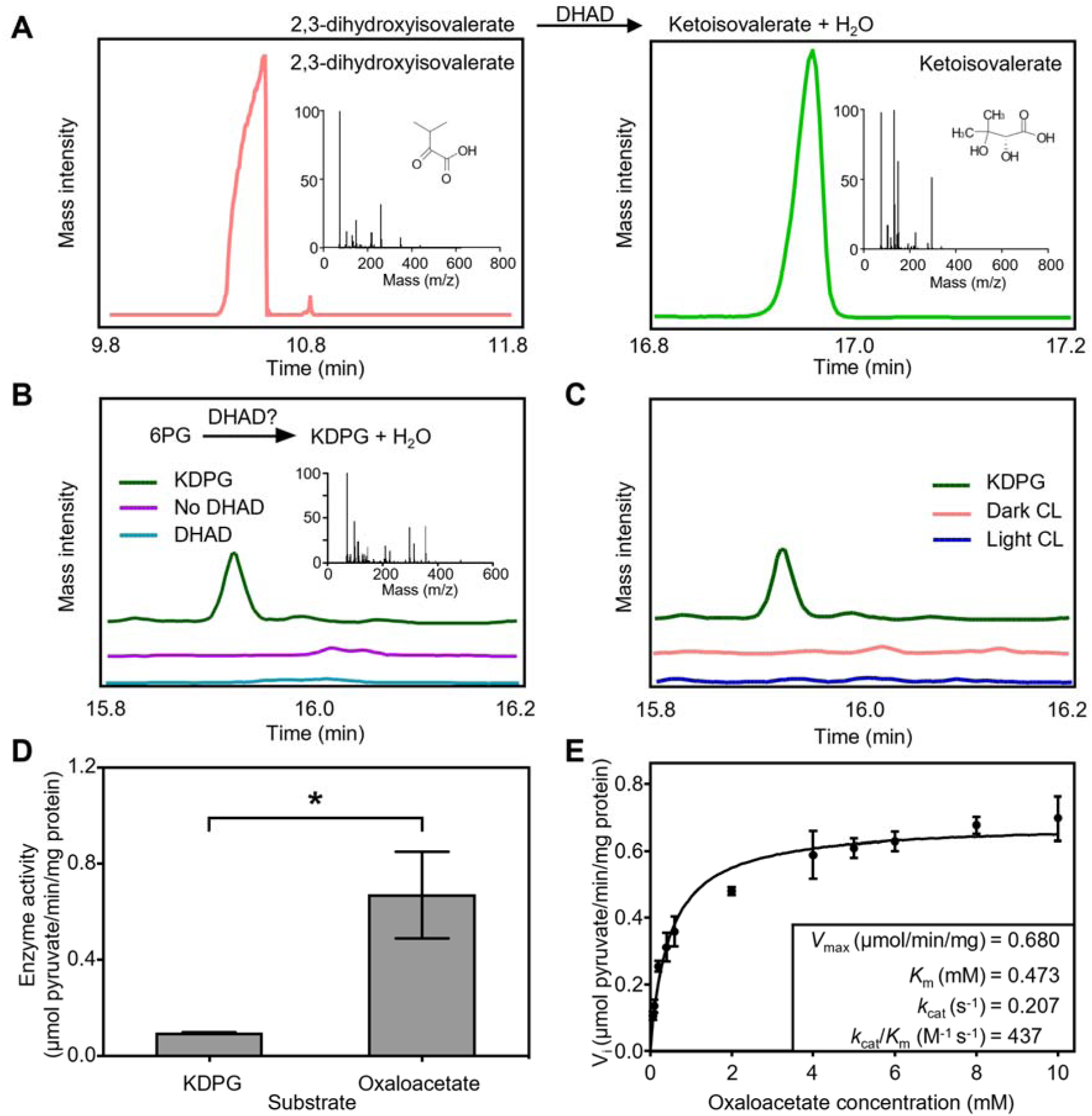
The ED pathway is incomplete in *S. elongatus*. **A)** DHAD from *S. elongatus* can catalyze the conversion of its native substrate 2,3-dihydroxyisovalerate into ketoisovalerate, detected by GC-MS. **B)** *S. elongatus* DHAD displays no apparent 6PG dehydratase activity. No KDPG was detected when purified DHAD was supplemented to the 6PG dehydratase enzyme assay (cyan). The purple line shows the no enzyme control. The green line shows the KDPG standard. **C)** 6PG dehydratase activity is not detected in the lysate of light (blue) or dark (pink) incubated cells. **D)** *S. elongatus* KDPG aldolase shows both KDPG cleavage activity and oxaloacetate decarboxylation activity. The asterisk indicates the higher activity on oxaloacetate than on KDPG. **E)** Kinetic analysis of KDPG aldolase using oxaloacetate as the substrate. The error bars in **D** and **E** represent the standard deviation of three independent measurements. Statistical significance was determined using a two-tailed Welch’s t-test. * *p* < 0.05

We further tested the activity of purified KDPG aldolase from *S. elongatus.* The result showed that *S. elongatus* KDPG aldolase can catalyze the conversion of KDPG to G3P and pyruvate (**Fig. 4D**). With an incomplete ED pathway in *S. elongatus*, it is likely that KDPG aldolase contributes to carbon metabolism through alternative activities. KDPG aldolase was reported to potentially catalyze several chemical reactions, one of which is oxaloacetate decarboxylation (Mavridis and Tulinsky, 1976). We thus tested whether *S. elongatus* KDPG aldolase can act on oxaloacetate. The *in vitro* assay showed that KDPG aldolase has decarboxylation activity on oxaloacetate to form pyruvate and CO_2_ (**Fig. 4D**), suggesting its potential involvement in carbon metabolism through this reaction in *S. elongatus*. The kinetics experiment further showed that the KDPG aldolase has a relatively low *k*cat/*K*m value of 437 M^-1^ s^-1^ for oxaloacetate (**Fig. 4E**). This seems to contradict the genetic result, where its deletion results in reduced biomass accumulation under diel cycle growth conditions (**Fig. 2B**). We reasoned that KDPG aldolase might be subject to unknown regulations that modulate its activity, which warrants further investigation. Collectively, the results support that KDPG aldolase, rather than the ED pathway, contributes to robust diel light/dark growth in *S. elongatus*.

### KDPG aldolase modulates carbon yield under diel cycles by conditionally bypassing TCA cycle activities in light

To explore the role of KDPG aldolase in modulating carbon yield under diel light/dark cycles, we sought to first identify contributing factors that affect its activity. The Δ*eda* mutant exhibited chlorosis under light conditions (**Fig. 2F**), indicating NADPH accumulation (*i.e.* a low ATP/NADPH ratio) and blockage of the photosynthetic electron transport chain (PETC). Additionally, while KDPG aldolase functions as a non-oxidative decarboxylase in *S. elongatus*, we hypothesized that NADP(H) could potentially serve as an allosteric regulator of the enzyme. We thus tested whether the dinucleotide could affect KDPG aldolase activities. Interestingly, incubation of purified KDPG aldolase with NADP^+^ but not NADPH led to an enhanced oxaloacetate decarboxylation activity (**Fig. 5A**). Further investigations using crude cell lysate showed no difference in oxaloacetate decarboxylation activity between the wild type and the Δ*eda* mutant when lysates alone were used, suggesting that KDPG aldolase is either in low abundance or inactive in wild type cells (**Fig. 5B**). However, pre-incubation of the lysates with NADP^+^ increased the oxaloacetate decarboxylation activity in the wild type relative to the Δ*eda* mutant (**Fig. 5B**), indicating that NADP^+^ activates KDPG aldolase. The results above suggest that KDPG aldolase could be activated when NADP^+^ levels exceed a certain threshold under diel light/dark cycles.

**Figure 5.**
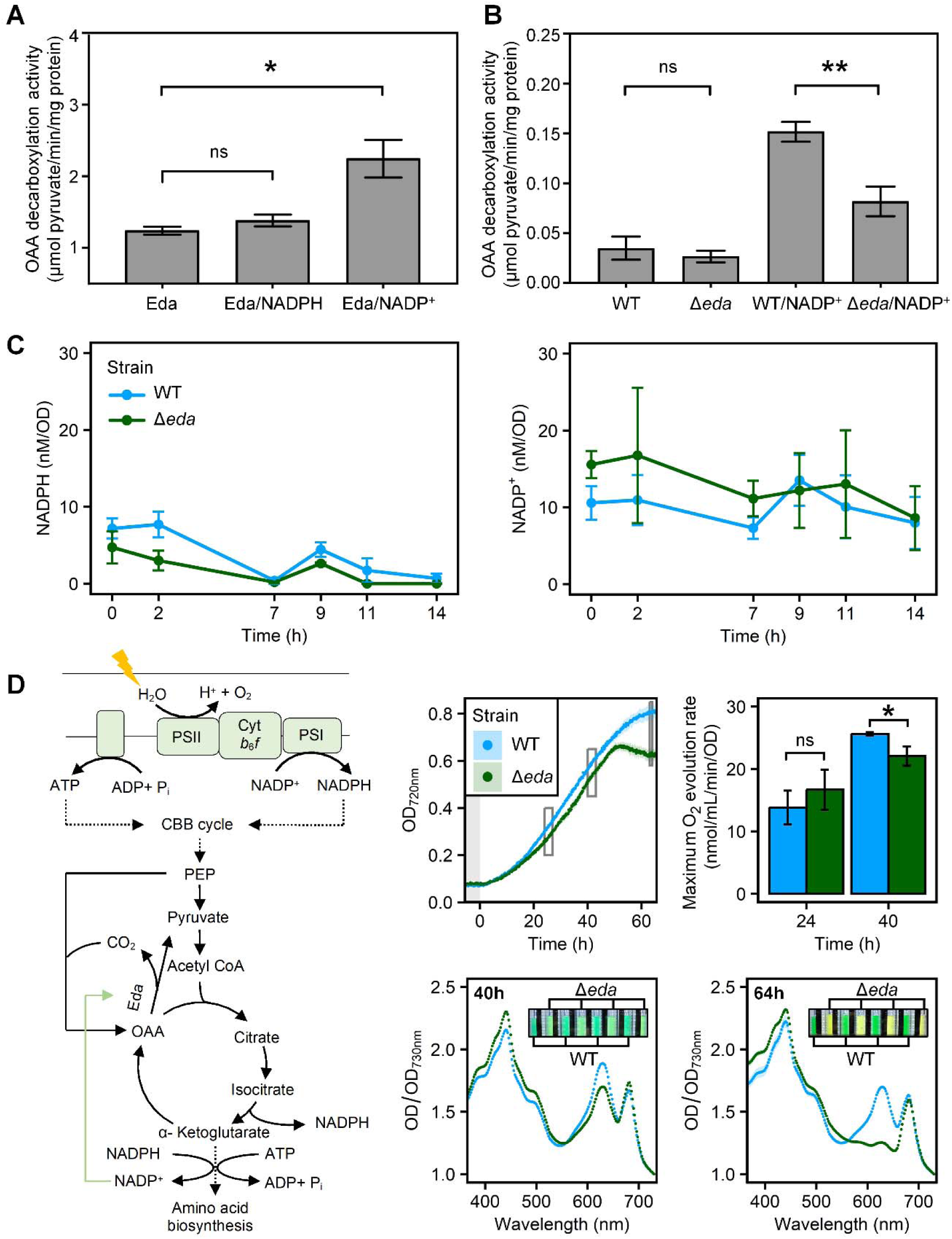
KDPG aldolase modulates carbon yield by fine-tuning TCA cycle activities in light. **A)** Supplementation with NADP^+^ increases the activity of purified KDPG aldolase (Eda) on oxaloacetate, whereas NADPH does not have this effect. **B)** Crude enzyme extracts from the wild-type strain exhibit higher oxaloacetate decarboxylation activity when supplemented with NADP^+^, compared to extracts from the Δ*eda* mutant. Without NADP^+^ supplementation, their activities are comparable. **C)** Levels of NADP(H) are monitored in both the wild type and the Δ*eda* mutant throughout the fourth dark phase of the 10h/14h light/dark cycles. **D)** A proposed model illustrates how KDPG aldolase modulates energy balance and carbon fixation in light. The Δ*eda* mutant exhibits growth defects, reduced oxygen evolution capacity, and a bleaching phenotype under prolonged continuous light. The specific time points for O_2_ evolution measurements or full visible light spectrum scanning are marked by squares on the growth curve. The error bars in panels **A** and **B** represent the standard deviation of three independent measurements. In panels **C** and **D**, the error bars or shading represent the standard deviation of independently cultivated biological quadruplicates. Statistical significance was determined using a two-tailed Welch’s t-test. * *p* < 0.05, ** *p* < 0.01, “ns” not significant

In cyanobacteria, the TCA cycle generates NADPH via oxidative decarboxylation reactions but does not produce ATP (Zhang and Bryant, 2011; Katayama et al., 2022). On the other hand, KDPG aldolase catalyzes the conversion of oxaloacetate to pyruvate, potentially bypassing TCA cycle activities in generating NADPH. We hypothesized that in the Δ*eda* mutant, the absence of KDPG aldolase leads to NADPH accumulation toward the end of the dark period in diel cycles. This accumulation could hinder electron transfer in the PETC and results in chlorosis upon light exposure. Following several cycles of light/dark growth, we thus measured the NADP(H) levels of both the wild type and the Δ*eda* mutant throughout the fourth dark period. Surprisingly, NADPH in both strains decreased to the lowest levels after the dark period, with a brief increase at the beginning of the latter half of the dark (**Fig. 5C**). In addition, both strains maintained similar and relatively stable NADP^+^ levels throughout the dark period (**Fig. 5C**). These results argue against our original hypothesis, instead suggesting that chlorosis in the Δ*eda* mutant is likely caused by relative NADPH accumulation in light.

Optimal photosynthetic carbon yield under diel cycles depends on balancing energy production from light reactions with energy consumption by carbon metabolism. When amino acid synthesis in light outpaces carbon fixation, it could lead to an imbalance in ATP/NADPH consumption relative to their production, thus reducing the PETC efficiency and exacerbating the shortage of carbon precursors needed for amino acid synthesis. α-ketoglutarate (AKG), a key intermediate in the TCA cycle, serves as the entry point for incorporating nitrogen into amino acids. Bypassing TCA activities thus allows for a temporary halt of amino acid synthesis and the rebalancing of ATP/NADPH production from light reactions. We propose that excessive NADPH consumption by amino acid synthesis could trigger a sharp rise of NADP^+^, thus activating KDPG aldolase to temporarily bypass TCA activities (**Fig. 5D**). To examine this possibility and minimize the influence of dark respiration, we cultivated cells under continuous light and monitored photosynthetic rates by measuring oxygen evolution. Supporting our hypothesis, the Δ*eda* mutant exhibited a lower photosynthetic rate than the wild type during the exponential growth phase. Furthermore, the Δ*eda* mutant showed severe chlorosis and ceased growing toward the later stage of the growth (**Fig. 5D**), underscoring the significant impact of KDPG aldolase on photosynthesis. Collectively, the results above strongly support the notion that KDPG aldolase optimizes daytime photosynthetic carbon yield by modulating TCA activities for amino acid synthesis.

### Phosphoketolase and KDPG aldolase mutants have disturbed cell metabolism under diel cycle growth

To further investigate how phosphoketolase and KDPG aldolase impact carbon yield under diel light/dark cycles, we conducted a comparative metabolomics study on *S. elongatus* wild type, Δ*pkt*, and Δ*eda* mutants grown under the 10h/14h light/dark cycle. Following a few cycles of light/dark growth, cells were collected in four different time points in the third dark period (**Supplementary Fig. S5**). Glycogen levels and degradation trends were similar among all three strains (**Fig. 6A**), indicating enough glycogen was synthesized in the light period even with the disruption of phosphoketolase or KDPG aldolase gene in the mutants.

**Figure 6.**
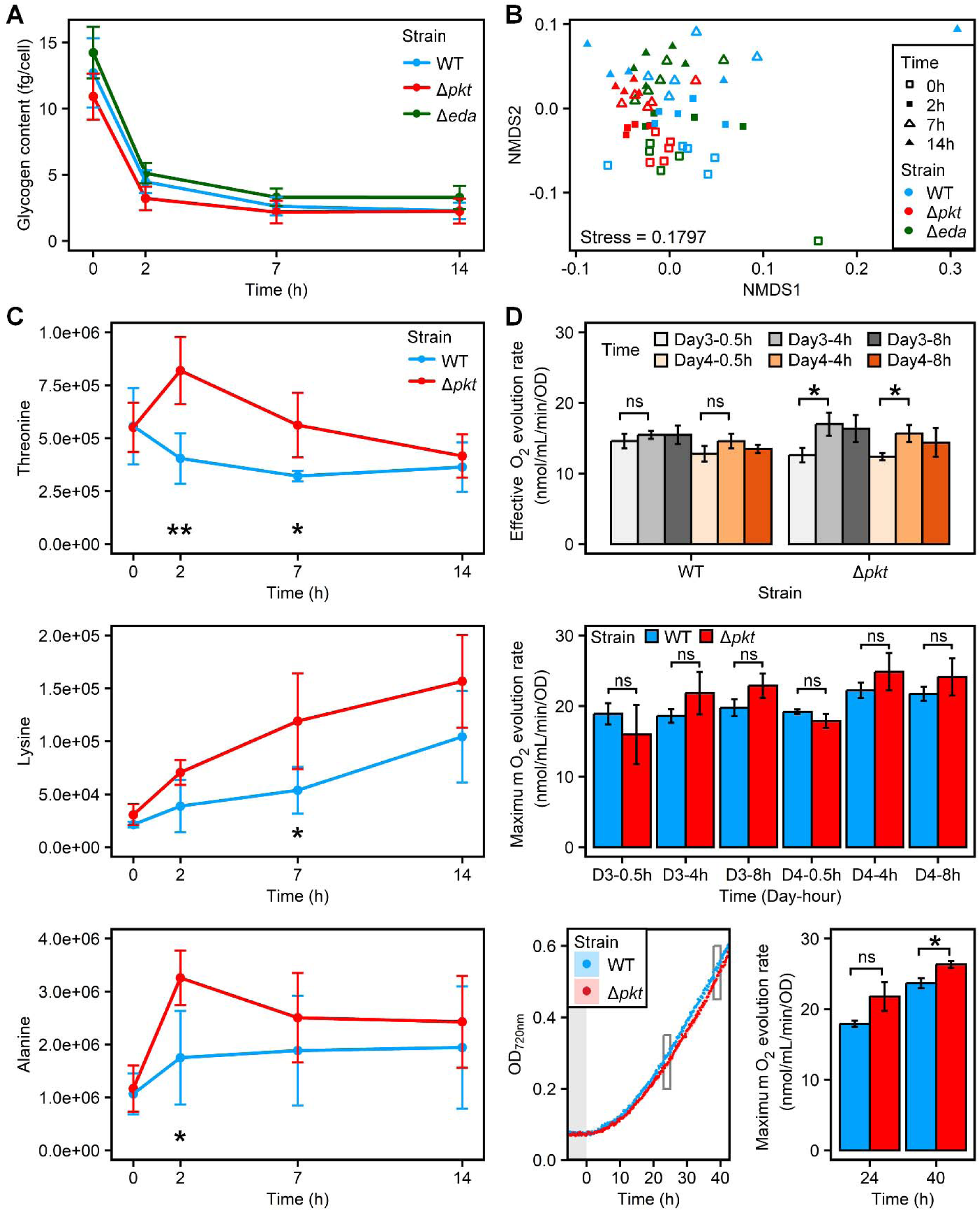
Phosphoketolase and KDPG aldolase mutants have disrupted metabolism in the dark. **A)** Glycogen degradation in the *S. elongatus* wild type, Δ*pkt*, and Δ*eda* mutants in the dark under 10h/14h light/dark cycle growth. **B)** A Bray-Curtis distance matrix highlights distinct metabolite profiles among the *S. elongatus* wild type, Δ*pkt*, and Δ*eda* mutants. **C)** Abundance changes of key intermediates in the dark suggest disrupted amino acid synthesis in the Δ*pkt* mutant. **D)** Oxygen evolution rates in the wild type and Δ*pkt* mutant grown under 10h/14h light/dark cycles (top and middle) and continuous light (bottom, with sampling time points indicated by squares on the growth curve). The error bars or shading represent the standard deviation of independently cultivated biological replicates (n = 5 for **A** and **C**; n = 3 for **D**). Statistical significance was determined using a two-tailed Welch’s t-test. * *p* < 0.05, ** *p* < 0.01, “ns” not significant

The Δ*pkt* and Δ*eda* mutants have disturbed cell metabolism compared to the wild type, indicated by distinct metabolite profiles in these strains; in each strain, the metabolite profiles also differ largely between the beginning of the night and latter periods of the dark (**Fig. 6B**; **Supplementary Table S2**). When looking at biomarkers that separate these strains, the Δ*pkt* mutant has the highest correlation with several amino acids (tyrosine, aspartate, threonine, and lysine), whereas the Δ*eda* mutant has the highest correlation with metabolites such as sarcosine, ribose-5-phosphate (R5P), oxoproline, and trans-4-hydroxyproline. The wild type, on the other hand, has the highest correlation scores with carbohydrates such as 2PG and 3PG (**Supplementary Table S3**). The disrupted metabolism in the Δ*eda* mutant compared to the wild type likely results from its inferior photosynthetic performance under light phases of the diel cycles.

Comparative metabolomics further suggests that phosphoketolase contributes to dark respiration by generating several precursors for amino acid synthesis. Phosphoketolase can convert xylulose-5-phosphate (X5P) to G3P and acetyl-phosphate (acetyl-P), the latter of which can be further converted to acetyl-CoA by acetate kinase and acetyl-CoA synthetase. Acetyl-CoA participates in amino acid synthesis by generating AKG through TCA reactions. If phosphoketolase contributes to significant acetyl-CoA flux, the lack of acetyl-P production by the Δ*pkt* mutant could result in a shortage of acetyl-CoA and further affect its reaction with oxaloacetate for AKG synthesis in the oxidative branch of the TCA pathway. This could lead to a transiently increased flux toward oxaloacetate and its downstream amino acids. Corroborating with this, we observed a surge of threonine in the first two hours of the dark and a consistently higher level of lysine in the Δ*pkt* mutant compared to the other two strains (**Fig. 6C**), both of which are derived from oxaloacetate. The lack of acetyl-CoA synthesis from acetyl-P likely needs to be compensated for by pyruvate decarboxylation, resulting in additional 3PG consumption for pyruvate generation. In support of this notion, we observed a higher increase of alanine, a derivative of pyruvate, in the Δ*pkt* mutant in the first two hours of the dark compared to the other two strains (**Fig. 6C**). These above results collectively show that phosphoketolase contributes to dark respiration by optimizing carbon flux toward generating precursors for amino acid synthesis.

The disrupted intermediate pools in the Δ*pkt* mutant likely affect its daytime photosynthesis, indicated by slower growth than the wild type during light phases of the diel cycles (**Fig. 2B**). We thus examined the daytime photosynthetic performance of both the wild type and the Δ*pkt* mutant grown under diel cycles. After two light/dark cycles, we measured their oxygen evolution rates at three time points during the light phases of day 3 and day 4 (0.5, 4, and 8 hours). On both days, maximum photosynthetic rates were comparable between these two strains. However, during the initial light period (0.5 hour), the Δ*pkt* mutant exhibited lower effective photosynthetic rates, measured using the growing light intensity, while the wild type quickly reached the maximum effective photosynthetic rate at 0.5 hour into the light phase (**Fig. 6D**). Further comparison of photosynthesis under continuous light conditions confirmed that the Δ*pkt* mutant had similar or even higher photosynthetic rates compared to the wild type under light conditions (**Fig. 6D**). These results strongly support that lower biomass yield in the Δ*pkt* mutant is primarily due to disturbed metabolism in the dark, which negatively impact photosynthesis when cells resume growth in light.

### Phosphoketolase and KDPG aldolase contribute to optimal photosynthetic carbon yield under diel light/dark cycles

To gain an overview of how phosphoketolase and KDPG aldolase contribute to robust diel growth, we conducted a time course proteomics on *S. elongatus* wild type under diel growth. The wild type was grown for three and a half cycles under 14h/10h light/dark cycles to obtain sufficient biomass, followed by sample collection at various time points of dark and light periods in the next 24 hours. In these proteomic samples, enzymes related to glycogen degradation (GlgP, glycogen phosphorylase) and the OPP pathway (Zwf, G6P dehydrogenase) were found in greater abundance in the dark than the light period (**Fig. 7**). This finding is consistent with the understanding of glycogen as the major carbon and energy source for dark respiration (Smith, 1983).

**Figure 7.**
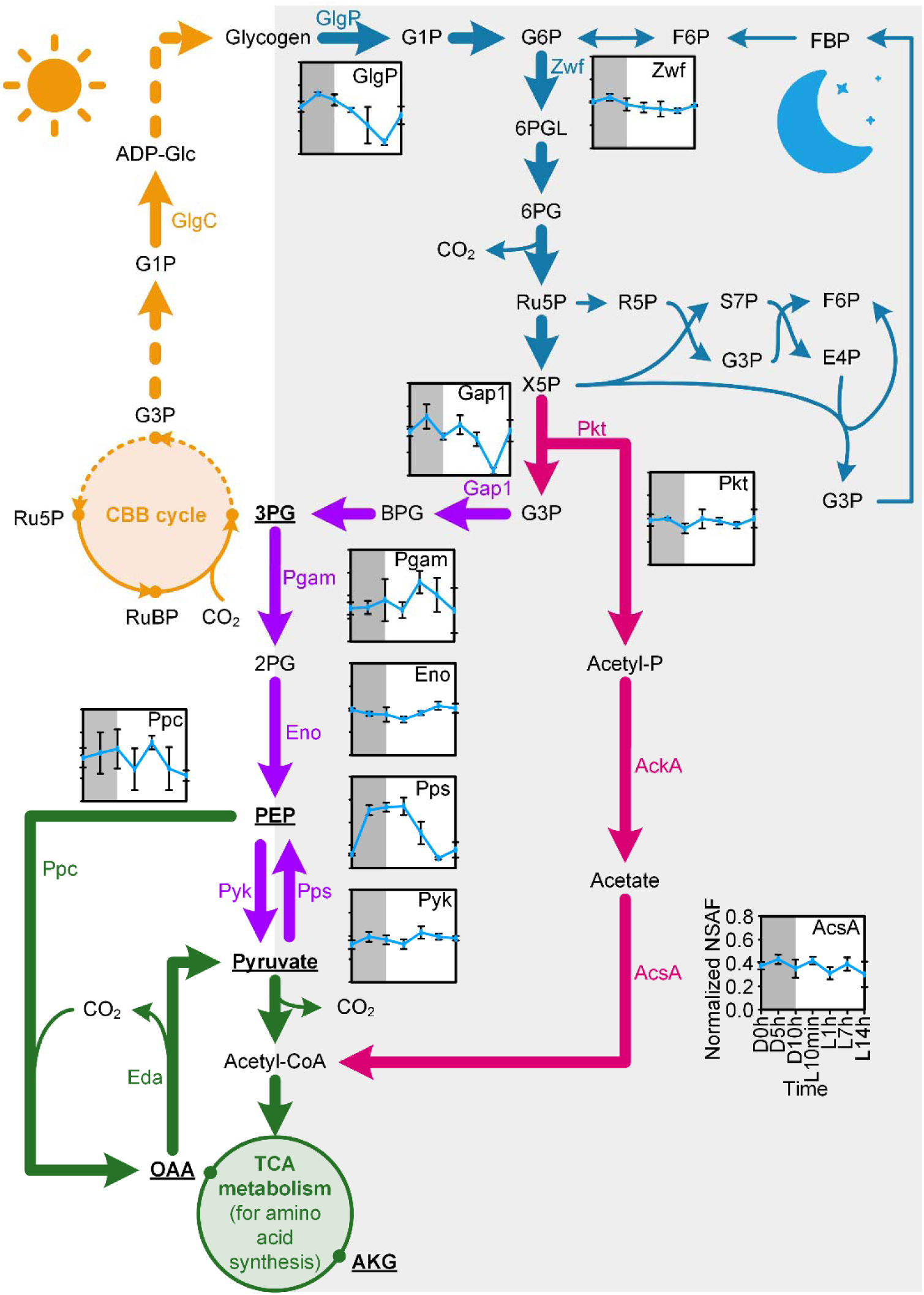
An integrative metabolic network ensures robust cyanobacterial diel growth. Gold arrows indicate the CBB cycle and glycogen synthesis during the light phase. Blue and rose red arrows represent the pentose phosphate pathway and phosphoketolase pathway in the dark, respectively. Purple arrows denote the downstream EMP pathway. Green arrows highlight metabolic processes contributing to amino acid synthesis. Metabolites labeled with bold fonts and underscores are precursors of amino acids. Inserted line plots show the dynamic changes in key enzymes involved in the 14h/10h light/dark cycle metabolism of *S. elongatus*. All inserted plots are uniformly scaled. The circle and error bar in each plot represent the mean and standard deviation of independently cultivated biological triplicates, respectively. Grey shading indicates the dark period of one diel cycle.

Next, phosphoketolase metabolism likely connects the OPP pathway with the downstream EMP pathway. Phosphoketolase directly converts C_5_ compounds derived from the OPP pathway, involving less enzymatic steps for G3P generation compared to the non-oxidative pentose phosphate (NOPP) pathway. In support of this notion, we found phosphoketolase and acetyl-CoA synthetase (AcsA) have consistently high abundance throughout the diel cycle (**Fig. 7**).

Furthermore, the downstream EMP pathway is involved in converging glycogen degradation products into key precursor metabolites for amino acid synthesis, including 3PG and phosphoenolpyruvate (PEP) (**Fig. 7**). G3P from the phosphoketolase reaction is first converted to 3PG in the downstream EMP pathway. Particularly, enzymes responsible for forming and oxidizing 3PG in the EMP pathway show distinct trends. On the one hand, G3P dehydrogenase (Gap1), the enzyme converting G3P to 1,3-bisphophoglycerate (BPG), increased in the first half of the night. On the other hand, phosphoglycerate mutase (Pgam) and enolase (Ena), responsible for converting 3PG to 2-phosphoglycerate (2PG) and PEP, were relatively stable in the dark (**Fig. 7**), potentially restricting 3PG conversion flux. The combined effort by these enzymes supports the notion that a portion of 3PG from glycogen degradation is diverted to amino acid synthesis.

Moreover, PEP synthase (Pps) increased to a significantly higher level in the first half of the night and stayed high throughout the dark, followed by a quick decrease to the base level in the next light period (**Fig. 7**), suggesting the conversion of pyruvate into PEP in the latter half of the night. This was further supported by the pyruvate kinase (Pyk) changes in the dark, with its level increasing during the first half of the night and decreasing afterwards (**Fig. 7**).

To synthesize amino acids derived from oxaloacetate and AKG, PEP carboxylase (Ppc) likely plays a crucial role in linking the downstream EMP pathway with the TCA cycle reactions. We found that Ppc, responsible for converting PEP to oxaloacetate, maintained at relatively high levels during the dark and in the middle of the light phase (**Fig. 7**), suggesting robust amino acids synthesis during these periods. Our results above underscore the importance of KDPG aldolase in balancing ATP/NADPH consumption relative to production for robust diel growth (**Fig. 5**). However, KDPG aldolase peptides were not detected in the wild-type proteomic samples, where an average of 1,000 proteins were identified per sample. This result aligns with the findings from the crude lysate experiments (**Fig. 5**), indicating that KDPG aldolase is likely present in low abundance and strictly activated by NADP^+^ to modulate TCA activities.

Collectively, the proteomics coupled with diel growth and biochemistry assays reveal an integrative respiratory pathway for dark respiration, where glycogen is first degraded to produce NADPH and C_5_ compounds via the OPP pathway, followed by conversion of C_5_ sugars to G3P and acetyl-P through the phosphoketolase metabolism. The intermediates G3P and acetyl-P further feed into downstream metabolism to generate precursors for amino acid synthesis, including 3PG, PEP, pyruvate, oxaloacetate, and AKG. Meanwhile, KDPG aldolase modulates TCA activities under light to ensure balanced ATP/NADPH production and consumption. The proposed route ensures optimal carbon and energy allocation in the dark for amino acid synthesis and balanced energy production and consumption during the light phase to support robust diel light/dark growth.

## Discussion

Respiration in the dark results in significant carbon loss and lowers the overall photosynthetic energy conversion efficiency in cyanobacteria and other photosynthetic organisms. Despite its importance, the strategies these organisms use to optimize carbon and energy allocation for dark respiration, and its contribution to photosynthesis, remain poorly understood. Here we show that in *S. elongatus*, cells form an integrative OPP-Phosphoketolase-EMP respiratory pathway in the dark to channel glycogen degradation products into key precursors for amino acid synthesis, which supports photosynthesis upon light exposure under diel light/dark cycles. We further show that the ED pathway in *S. elongatus* is incomplete, with KDPG aldolase contributing to robust diel growth through an alternative decarboxylation activity on oxaloacetate.

Phosphoketolase is an important glycolytic enzyme in many bacteria but its role in cyanobacteria is under debate (Årsköld et al., 2008). Previously, phosphoketolase was shown to carry significant carbon flux under mixotrophic conditions in an engineered *Synechocystis* sp. PCC 6803 strain (Δ*glgC/xylAB*), which lacks the ability to synthesize glycogen but can utilize xylose (Xiong et al., 2015). We show here that phosphoketolase metabolism is critical for dark respiration under light/dark diel growth in the obligate photoautotrophic *S. elongatus*. When cyanobacteria enter the dark, phosphoketolase reaction connects the OPP pathway with amino acid synthesis for cell maintenance in the dark (**Fig. 7**). A recent bioinformatics study analyzed the co-occurrence of key enzymes from four glycolytic pathways (EMP, ED, OPP, and phosphoketolase) in cyanobacteria and found that enzymes from the phosphoketolase and OPP pathways co-occur with the highest probability of 84.8% (Bachhar and Jablonsky, 2022), supporting the OPP-phosphoketolase route in the dark during cyanobacterial diel light/dark growth. In addition, phosphoketolase exists in eukaryotic phototrophs such as diatoms (*Chaetoceros debilis*, *Leptocylindrus danicus*, *Amphora coffeiformis*, and *Phaeodactylum tricornutum*), haptophytes (*Diacronema lutheri*), dinoflagellates (*Symbiodinium microadriaticum*), and some land plants (*Capsicum annuum*, *Salix suchowensis*, and *Gossypium arboreum*) (**Supplementary Fig. S6**), suggesting its role in a wide range of photosynthetic organisms.

Phosphoketolase was found to be expressed at consistent levels in the wild type throughout a diel light/dark cycle (**Fig. 7**). If phosphoketolase is functioning under light, its activity toward either X5P or F6P would likely lead to unstable CBB cycle reactions and retarded growth (Yu et al., 2018). A previous research showed that phosphoketolase has much higher activities on X5P than F6P in *S. elongatus*, with both activities significantly inhibited by ATP (Chuang and Liao, 2021). This helps explain the similar growth of the wild type and the Δ*pkt* mutant under continuous light (**Fig. 2D, 6D**), suggesting that phosphoketolase activity is likely regulated mainly at the allosteric rather than transcriptional level during diel growth. When ATP level lowers upon cell entering the dark (Rust et al., 2011), phosphoketolase can be activated to catalyze X5P cleavage. A more recent report confirmed the allosteric regulation of phosphoketolase by ATP, and further showed an increased phosphoketolase activity during light-dark transitions to brake on the CBB cycle and switch cell metabolism into respiration (Lu et al., 2023).

Phosphoketolase metabolism further contributes to robust diel cell growth by minimizing carbon loss in the dark and supporting optimal photosynthetic growth upon light exposure. When phosphoketolase was deleted in the Δ*pkt* mutant, lower biomass yield was observed under the 10h/14h light/dark cycles (**Fig. 2B; Supplementary Fig. S7**). Without phosphoketolase, the Δ*pkt* mutant will likely obtain G3P through transketolase reactions in the NOPP pathway by converting R5P and X5P into G3P and sedoheptulose-7-phosphate (S7P) or converting erythrose-4-phosphate (E4P) and X5P into G3P and fructose 6-phosphate (F6P) (**Fig. 7**). The siphoning of G3P from NOPP leads to imbalanced carbon pools, likely resulting in S7P and F6P accumulation. Although we did not detect these phosphorylated sugars in our metabolomics, this hypothesis is supported by findings in a previous study where *S. elongatus* Δ*pkt* was grown to high density and placed in darkness. In this previous study, higher levels of S7P and hexose-6P were found in the Δ*pkt* mutant than the wild type cells in the dark (Chuang and Liao, 2021). The imbalanced intermediate pools in the Δ*pkt* mutant could adversely affect daytime photosynthesis, potentially explaining the observed lower photosynthetic efficiency during the initial light phase of diel growth (**Fig. 6D**). In addition, lower amount of 3PG, PEP, and acetyl-CoA were found in the Δ*pkt* mutant than the wild type, whereas pyruvate was accumulated to a higher level in the Δ*pkt* mutant (Chuang and Liao, 2021). This is likely due to increased consumption of 3PG in the Δ*pkt* mutant to generate pyruvate and acetyl-CoA in the downstream EMP pathway, which would lead to higher carbon loss in the pyruvate decarboxylase reaction. This suggests phosphoketolase metabolism contributes to significant flux for acetyl-CoA generation from acetyl-P, explaining the observed acetyl-CoA accumulation in the wild type in the previous study (Chuang and Liao, 2021). However, this hypothesis needs to be tested by verifying activities of the two enzymes involved in converting acetyl-P to acetyl-CoA in the dark.

KDPG aldolase contributes greatly to the robust diel growth in *S. elongatus*, as shown by much lower biomass accumulation in the Δ*eda* mutant compared to the wild type under light/dark cycles (**Fig. 2B**). A similar retarded growth was observed in the Δ*eda* mutant of *Synechocystis* sp. PCC6803 when cells grew photoautotrophically under light/dark cycles (Chen et al., 2016). This previous study attributed the growth defect to the absence of the ED pathway. However, no active 6PGDH activity or KDPG metabolite were detected in *S. elongatus* (**Fig. 4**). In addition, the *in vitro* enzyme assay of KDPG aldolase shows a much higher activity toward oxaloacetate than KDPG (**Fig. 4D**). Our results thus strongly support that the ED pathway is incomplete in *S. elongatus* with the KDPG aldolase supporting robust diel growth through its alternative decarboxylation activity. This is also consistent with the findings that the KDPG aldolase is much more prevalent than 6PGDH in cyanobacteria (Bachhar and Jablonsky, 2022). Recent evidence further points to the absence of the ED pathway in plant ancestors due to the lack of genuine 6PGDH in their organisms (Evans et al., 2024), making it particularly intriguing to clarify the role of KDPG aldolase in photosynthetic organisms.

The significance of KDPG aldolase is demonstrated by its role in oxaloacetate metabolism. Oxaloacetate connects the downstream EMP pathway with the TCA cycle, which is responsible for generating AKG, a key precursor for incorporating nitrogen into all amino acids (**Fig. 5D**). Under light conditions, excessive amino acid synthesis can disrupt the balance of ATP/NADPH consumption relative to its production by light reactions, potentially resulting in a lower ATP/NADPH ratio and reduced photosynthetic efficiency. KDPG aldolase plays a critical role in modulating this balance by temporarily bypassing the TCA cycle through oxaloacetate decarboxylation. This regulation is essential for optimal photosynthetic carbon yield by aligning amino acid synthesis with energy production from light reactions. This necessitates a precise control mechanism that limits the bypass of the TCA cycle to conditions of high NADP^+^ levels, which are indicative of excessive synthesis activities in cells. This stringent regulation is maintained through the low expression and enzyme kinetics of KDPG aldolase (**Fig. 4E**), which is activated only when NADP^+^ levels exceed a specific threshold. However, the detailed regulation mechanisms require further investigation to be fully understood. Additionally, this regulation is not required for amino acid synthesis in the dark, as ATP/NADPH production is not coupled by light reactions, and respiration instead is responsible for ATP generation. This is supported by our observations that NADP(H) changes are comparable between the wild type and the Δ*eda* mutant (**Fig. 5C**).

Overall, our study illustrates how phosphoketolase and KDPG aldolase contribute to optimizing photosynthetic carbon yield under diel light/dark cycles in cyanobacteria. Further validation of the proposed roles in algae and plants could help understand the prevalence of these enzymes in phylogenetically diverse phototrophic taxa and elucidate how compartmentation in eukaryotes alters the role of glycolytic routes in dark respiration and the robustness of photosynthesis.

## Methods

### Strain construction

Plasmids required for generating mutant strains were prepared using HiFi DNA assembly (NEB, USA) and then transformed into *S. elongatus* wild type. The DNA fragments for construction of the plasmids were obtained through either high-fidelity amplification using Q5 DNA Polymerase (NEB, USA) or restriction enzyme digestion. The strains and plasmids information can be found in **Supplementary Fig. S1** and **Table S1**. The mutant strains were confirmed by segregation PCR (**Supplementary Fig. S2**).

### Growth characterization

The wild type and mutant strains of *S. elongatus* were grown in BG11 medium supplemented with 20 mM NaHCO_3_ and 10 mM N-[Tris(hydroxymethyl)methyl]-2-aminoethanesulfonic acid (TES, pH 7.8) at 30 °C under 100 µmol/m^2^/s (otherwise specified) cool white LED illumination or different diel cycles in the Multi-Cultivator MC 1000-OD (Photon Systems Instruments, Czech Republic). Growth was directly monitored by recording the absorbance at 720 nm at 10 min intervals using the built-in optical density sensor. For the mutant strains, appropriate antibiotics were supplemented to the growth medium (5 µg/mL kanamycin for Δ*glgC*; 2 µg/mL gentamicin for Δ*pfk*; 5 µg/mL chloramphenicol for Δ*pkt*; 2 µg/mL spectinomycin and 2 µg/mL streptomycin for Δ*eda*; 5 µg/mL chloramphenicol, 2 µg/mL spectinomycin, and 2 µg/mL streptomycin for Δ*pkt*Δ*eda*). Cultures for proteomic and metabolomic analyses were grown in the same medium and light conditions as described above.

### Glycogen quantification

The glycogen content was quantified to compare its degradation trends in the dark between the wild-type and mutant strains, following the modified protocol as described previously (De Porcellinis et al., 2017). For each sample, 5 mL of cyanobacterial cells were collected and centrifuged at 9000 g and 4 °C for 20 min, washed with 1 mL of ice-cold Tris-HCl buffer (50 mM, pH 8.0), and stored at -30 °C as cell pellets until extraction of glycogen. To extract the glycogen, the pellet samples were kept on ice, resuspended in 500 μL of the cold Tris-HCl buffer, and lysed by bead beating for 20 cycles of 20 sec on (at 3800 rpm) and 2 min off (on ice). The cell lysate was collected by centrifugation at 9,000×g and 4 °C for 20 min, 150 μL of which was mixed with 1350 μL of 96% ethanol and incubated at 90 °C for 10 min to remove chlorophyll. Then, the samples were incubated on ice for 30 min and centrifuged at 20,000 g at 4 °C for 30 min to precipitate the glycogen. The resulting pellet was air-dried for 15 min at room temperature and resuspended in 200 μL of 50 mM sodium acetate buffer (pH 5.0) which contained 8 U/mL amyloglucosidase and 2 U/mL α-amylase. The mixture was vortexed vigorously and incubated at 60 °C for 2 h to hydrolyze the glycogen into glucose. The samples were centrifuged at 10,000×g for 5 min, and the supernatant was collected to quantify the glucose concentration using glucose oxidase peroxidase (GOD-POD) assay (Megazyme, USA) in the SpectraMax iD5 microplate reader (Molecular Devices). The glycogen content was normalized by the corresponding cell number, which was determined by an Attune NxT Flow Cytometer (Invitrogen).

### Determination of dark respiration rates and oxygen evolution rates

To measure the dark respiration rates of *S. elongatus* wild type and mutant strains under the 10h/14h light/dark diel growth, six independently cultivated biological replicates (triplicates from two independent experiments) of each strain were collected from the PSI multi-cultivator at the end of the fourth light period. The collected cultures were aliquoted into 2-mL tubes to ensure efficient reoxygenation from the air, and incubated in a rotator at 30°C in the dark for 0, 2, 9, 11, and 14 hours until measuring the respiration rates using the Oxytherm+P system (Hansatech Instruments) at 30°C. The respiration rates were normalized by OD_720nm_ (recorded by the PSI multi-cultivator).

To measure the oxygen evolution rates of *S. elongatus* wild type and mutant strains under the continuous light or under the light phase of light/dark cycles, 2 mL culture was sampled from the PSI multi-cultivator and immediately transferred into the chamber of the Oxytherm+P system (Hansatech Instruments). The oxygen evolution rates were measured at 30 °C, with the maximum rates measured under the saturated light and the effective rates measured under a light about 100 µmol/m^2^/s. The oxygen evolution rates were corrected by the respiration rates and normalized by OD_720nm_ (recorded by the PSI multi-cultivator).

### Derivatization and detection of KDPG and other metabolites by gas chromatography-mass spectrometry (GC-MS)

The samples and standards used in GC-MS analysis were prepared using the N-methyl trifluoroacetamide (MSTFA) derivatization method. For derivatization reactions, 40 μL of 2% methoxyamine hydrochloride (dissolved in pyridine) was added to freeze-dried samples, followed by shaking at 37 °C for 2 h. After a brief spin, 70 μL of MSTFA with 1% trimethylchlorosilane (TMCS) was added, followed by shaking at 37 °C for 30 min. After centrifugation at 17,000 g for 5 min, about 80 μL of the clear supernatant was transferred into a GC vial. KDPG standards spanning from 11 to 176 pmol, supplemented with L-norvaline as the internal standard, were prepared, dried, and derivatized in parallel with the samples to establish a calibration curve for KDPG detection. The derivatized samples were run on a Thermo GC-MS equipment coupled with the Zebron ZB-5MSplus column. The temperature program was set to 85 °C with an initial hold for 2 min, followed by 10 °C/min of increase till 110 °C, 5 °C/min of increase till 230 °C, and 15 °C/min of increase till 320 °C, with a final hold for 2 min. The MS scan was set for the range of 70-600 amu.

### Crude enzyme assays for 6PG dehydratase activity

Crude enzyme assay was used to determine potential 6PG dehydratase activity by *S. elongatus*. For crude protein extraction, *S. elongatus* wild type cells were grown in liquid BG11 medium until OD_730nm_ reached 2. After 6 hours of dark adaptation, 10 mL of each culture was centrifuged at 8,000×g for 10 minutes at 4 °C. The cell pellet was washed twice with 5 mL of cold 50 mM Tris-HCl buffer (pH 8) containing 1× Protease Inhibitor Cocktail, and finally resuspended in 1000 µL of this buffer. The resuspended cells were lysed using 0.1 mm diameter silicone beads in a bead beater for 15 cycles of 10 sec on and 2 min off on ice. The supernatant containing the crude protein was separated by centrifugation at maximum speed for 10 minutes at 4°C. The protein concentration in the supernatant was determined by Bradford assay.

The crude lysate reaction mix containing 50 mM Tris-HCl buffer, 100 mM NaCl and 100 µL of the crude lysate was preincubated at 30 °C for 15 minutes. The reaction was initiated using 2.5 mM 6-phosphogluconic acid as the substrate and then incubated at 30 °C for two hours under both light and dark conditions. Follow incubation, samples were analyzed for KDPG production using GC-MS.

### Recombinant enzyme purification and *in vitro* enzyme assays

To check the activity of DHAD (*ilvD*) and KDPG aldolase (*eda*), the gene fragment was amplified from the genome of *S. elongatus*, linked with the His-tag containing pET28A vector using Gibson assembly, and transformed into *E. coli* BL21 competent cells. To extract the his-tagged protein, 100 mL of the transformant cells were grown until OD_600nm_ reached ∼ 0.5, when 0.1 M IPTG was added to induce the overexpression of the His-tagged protein. After 4 h of induction, the culture was centrifugated at 6,000 rpm and 4 °C for 10 min. The cells were washed twice with 5 mL of cold 100 mM Tris-HCl buffer (pH 8), resuspended in 1 mL of the buffer supplemented with 1× Protease Inhibitor Cocktail, and then lysed by 0.1 mm diameter silicone beads in a bead beater for 15 cycles of 10 sec on and 30 sec off on ice. The lysate was collected after centrifugation at maximum speed for 10 minutes at 4 °C. The targeted His-tagged protein was purified using the NEB Express Ni Spin Columns (catalog no. S1427) following the manufacturer’s protocol and confirmed by SDS-PAGE **(Supplementary Fig. S8)**.

To test KDPG aldolase activities, the His-tagged KDPG aldolase enzyme was coupled with Lactate Dehydrogenase (LDH): NADH assay to detect pyruvate generation using KDPG or oxaloacetate as the substrate. The reaction mixture contained 100 mM Tris-buffer (pH 8), 150 mM NaCl, 2 mM NADH, 6 units of LDH, 5 µL of purified KDPG aldolase, and 2 mM of the required substrate. The rection mixture without substrate was preincubated for 15 minutes at 30 °C and later 2 mM substrate was added to initiate the reaction. The reactions without substrate and without enzyme were prepared separately as controls. The initial absorbance (OD_340nm_) was read in the microplate reader and the incubation was maintained until a drop in the absorbance was observed. The absorbance at 340 nm was read every 10 minutes to generate NADH oxidation curve over time.

To determine whether DHAD can function as 6PGDH to catalyze the conversion of 6PG to KDPG, the reaction mixture containing 50 mM Tris HCl (pH 7.8), 150 mM NaCl, 1 mM MgCl_2_ and 10 µL purified DHAD was preincubated for 15 minutes. The reaction was initiated using 2.5 mM 6PG as substrate and was incubated at 30 °C for 1 hour. The reaction was then anaylzed using GC-MS for production of KDPG as the product. The native DHAD activity for conversion of 2,3-dihydroxyisovalerate to ketoisovalerate was used as positive control. For this activity, the reaction was initiated using 2,3-dihydroxyisovalerate as the substrate and production of ketoisovalerate was analyzed using GC-MS.

### Enzyme kinetics of KDPG aldolase on oxaloacetate

The oxaloacetate decarboxylation activity of KDPG aldolase was measured at different substrate concentrations to generate the Michaelis-Menten curve for the calculation of *K*_m_ and V_max_ values. The reaction mix containing 100 mM Tris buffer (pH 8), 150 mM NaCl, 2 mM NADH, 6 units of LDH and 5 µL of purified KDPG aldolase was preincubated for 15 minutes at 30 °C. The reaction was initiated using a range of substrate concentrations (0.075-10 mM) and was allowed to incubate at 30°C for 20 minutes. The drop in OD_340nm_ was normalized to the no-enzyme control due to spontaneous decarboxylation of oxaloacetate. The experimental data for the initial slope was used for kinetics measurements and was fitted through the Michaelis-Menten equation using Graphpad software for calculation of *V*_max_, *K*_m_, and *k*_cat_ values.

### Effects of NADP(H) on KDPG aldolase activity

To test the effect of NADP^+^ and NDAPH on the oxaloacetate decarboxylation activity of KDPG aldolase, 1 mM NADP^+^ or NADPH was added to the reaction mix containing PBS buffer (pH 7.4), 2 mM NADH, 6 units of LDH and 1 μg of purified KDPG aldolase. The reaction mix with NADP^+^ or NADPH was pre-incubated at 30°C for one hour. After this, the reaction was initiated by adding 2 mM oxaloacetate as the substrate. The reactions without substrate and without enzyme were prepared separately as controls. The initial absorbance (OD_340nm_) was read in the microplate reader and the incubation was maintained until a drop in the absorbance was observed. The drop in OD_340nm_ was normalized to the no-enzyme control due to spontaneous decarboxylation of oxaloacetate.

To test KDPG aldolase activity in *S. elongatus* cell lysates and the effect of NADP^+^, both wild type and Δ*eda* mutant were grown to log phase for crude protein extraction. The reaction mixtures, both with and without 1 mM NADP^+^, contained PBS (pH 7.4), 3 units of LDH, and 25 µg of crude protein lysate. After a 30-minute pre-incubation, 2 mM oxaloacetate and 2 mM NADH were added. The absorbance (OD_340nm_) was read in the microplate reader and the incubation was maintained until a drop in the absorbance was observed. The drop in OD_340nm_ was normalized to the no-enzyme control due to spontaneous decarboxylation of oxaloacetate.

### Measurement of NADP(H) levels

Quadruplicates of *S. elongatus* wild-type and mutant strains were cultivated under 10h/14h light/dark cycles. At specified times (0h, 2h, 7h, 9h, 11h, and 14h) during the fourth cycle’s dark period, 10 mL of cells were harvested, quenched with ice-cold PBS slurry (pH 7.4), and centrifuged to collect cell pellets, which were stored at -80°C. For NADP(H) extraction, pellets were resuspended in 0.5 mL of ice-cold PBS and lysed with 0.5 mL of 0.2N NaOH containing 1% dodecyltrimethylammonium bromide. After centrifugation at maximum speed for 20 min at 4 °C, the supernatant was collected to determine the NADP^+^ and NADPH concentrations using Promega NADP/NADPH-Glo Assay (catalog no. G9081). The amounts of NADP^+^ and NADPH in each sample were calculated from the standard curves and normalized to OD_720nm_ (recorded by the PSI multi-cultivator).

### Metabolomics

Five independently cultivated biological replicates of Δ*pkt*, Δ*eda*, and *S. elongatus* wild type strains were sampled at 0h, 2h, 7h, and 14h of the dark period of the third 10h/14h light/dark cycle, following a modified protocol described previously (Wang and Young, 2022). For each sample, 10 mL of cyanobacterial culture (10^8^∼10^9^ cells) was filtered through a 0.45 μm nylon filter by vacuum filtration. The resulting filter was immediately inverted into 1.7 mL of precooled (-80 °C) methanol in the 6-well plate sitting on dry ice. After incubation at -80 °C for a few hours, the cells were scraped off from the filter, and the mixture of cells and methanol was transferred into a 2-mL centrifuge tube and dried by SpeedVac. The dried samples were stored at -80 °C until shipping to the West Coast Metabolomics Center for primary metabolism analysis on a GC-Time of Flight (TOF) mass spectrometer. A small aliquot (about 0.5 mL) of the culture was saved before filtration for flow cytometry analysis to estimate the cell number in each sample, which was later used to normalize the metabolite levels. The glycogen contents were also measured for each sample.

### Shotgun proteomics

Time-course proteomics were conducted for *S. elongatus* wild type during a 14h/10h light/dark cycle after a few growth cycles. About 4∼6 OD of cells were collected in biological triplicates at 0 h, 5 h, and 10 h of the dark period and at 10 min, 1 h, 7 h, and 14 h of the following light period. The cells were quenched and washed by ice-cold PBS slurry (pH 7.4), centrifugated at maximum speed to obtain the cell pellets, and stored at -30 °C until protein extraction. To lyse the protein, the cell pellets were resuspended in 500 μL of lysis buffer comprising 50 mM Tris-HCl (pH 8.0), 0.02 % n-Dodecyl-Beta-Maltoside, and 1× Protease Inhibitor Cocktail, and homogenized by bead beating for 10 cycles of 15 sec on at 3800 rpm and 2 min off on ice. After centrifugation at 20,000×g and 4 °C for 30 min, the supernatant was collected for protein concentration measurement using the Bradford assay (Thermo Scientific) and for downstream protein digestion.

For each sample, 100 μg of total protein was incubated with 8 M urea and 5 mM dithiothreitol (DTT) at 37 °C for 1 h, and then with 15 mM iodoacetic acid (IAA) at room temperature in dark for 30 min, followed by addition of 50 mM Tris-HCl (pH 8.0) to reduce the urea concentration to 2 M. The 100 μg of protein in each diluted sample was digested by 2 μg of Trypsin Gold (Promega, USA) at 37 °C for 20 h. The resulting peptides were purified by the Sep-Pak C18 column (Waters) and separated by the Pierce High pH Reverse-Phase Peptide Fractionation Kit (Thermo Scientific) into 8 fractions, which were further run on an EASY-nLC 1000 chromatography coupled to a LTQ Orbitrap XL mass spectrometer (Thermo Scientific, USA) for LC-MS/MS analysis. Raw MS/MS data were searched and analyzed using the PatternLab V software (Carvalho et al., 2016). The normalized spectral abundance factor was used to compare protein expression changes across the diel cycle. The mass spectrometry proteomics data can be located under the dataset identifier MSV000093712 in the MassIVE repository or with the dataset identifier PXD048049 under the ProteomeXchange Consortium (Vizcaíno et al., 2015).

### Phylogenetic analysis

Phosphoketolase protein sequences were searched in the UniProt database. Two representative sequences for cyanobacteria and available sequences for eukaryotic photoautotrophs (dinoflagellates, diatoms, haptophytes, and Viridiplantae) were downloaded for phylogenetic analysis. A maximum likelihood tree was constructed in MEGA 7, using the automatically selected best-fit substitution model, after the sequences were aligned with MUSCLE and curated manually (Kumar et al., 2016).

### Statistical analysis

The Welch’s t-tests were performed to compare the differences in growth (OD_720nm_), pigment absorbances, oxygen evolution and respiration rates, metabolite abundances, and enzyme activities between different strains, time points, or treatments. The terminal OD_720nm_ under the light/dark cycles is defined as the average OD_720nm_ in the last hour of the last light period. The terminal OD_720nm_ under the continuous light is defined as the average OD_720nm_ of the last hour. Two-tailed t-tests were used unless otherwise specified.

Pairwise comparisons in metabolomic profiles between different strains and time points were performed by permutation MANOVAs based on the Bray-Curtis distance matrix, after confirming equivalent beta-dispersion. *p*-values were adjusted for false discovery rate (FDR). The similarities or dissimilarities across all samples in their metabolomic profiles were visualized using Non-metric Multidimensional Scaling (NMDS) based on the Bray-Curtis distance matrix. The coefficients of the top 10 metabolites associated with each strain in the dark period (2h, 7h, and 14h) were determined using Partial Least Squares Discriminant Analysis (PLS-DA).

## Acknowledgements

We thank Julie Maupin-Furlow for insightful suggestions and careful reading of the manuscript. We thank Rachael Morgan-Kiss for providing access to the PSI multi-cultivator and the Center for Bioinformatics and Functional Genomics at Miami University for providing access to several pieces of equipment used in this study. This work was supported by the National Science Foundation grant number 2414925 (Original award number 2042182). X.W. would like to acknowledge support from both University of Florida and Miami University.

## Author contributions

X.W. conceived the project. N.X. performed the genetics, proteomics, growth physiology, respiration, and photosynthesis measurement experiments with assistance from K.R. C.S. performed the enzyme experiments related to KDPG aldolase. N.X. and X.W. wrote the manuscript with edits from C.S.

## Data availability

All data are presented in the main text and supplementary materials. Proteomics data has been deposited into the public database MassIVE repository and ProteomeXchange Consortium.

## References

Amthor, J.S., Bar-Even, A., Hanson, A.D., Millar, A.H., Stitt, M., Sweetlove, L.J., and Tyerman, S.D. (2019). Engineering strategies to boost crop productivity by cutting respiratory carbon loss. Plant Cell 31, 297–314.

Årsköld, E., Lohmeler-Vogel, E., Cao, R., Roos, S., Rådström, P., and van Niel, E.W.J. (2008). Phosphoketolase pathway dominates in ATCC 55730 containing dual pathways for glycolysis. J Bacteriol 190, 206–212.

Bachhar, A., and Jablonsky, J. (2022). Entner-Doudoroff pathway in *Synechocystis* PCC 6803: Proposed regulatory roles and enzyme multifunctionalities. Front Microbiol 13, 967545.

Carvalho, P.C., Lima, D.B., Leprevost, F.V., Santos, M.D., Fischer, J.S., Aquino, P.F., Moresco, J.J., Yates III, J.R., and Barbosa, V.C. (2016). Integrated analysis of shotgun proteomic data with PatternLab for proteomics 4.0. Nat Protoc 11, 102.

Chen, X., Schreiber, K., Appel, J., Makowka, A., Fahnrich, B., Roettger, M., Hajirezaei, M.R., Sonnichsen, F.D., Schonheit, P., Martin, W.F., and Gutekunst, K. (2016). The Entner-Doudoroff pathway is an overlooked glycolytic route in cyanobacteria and plants. Proc Natl Acad Sci USA 113, 5441–5446.

Chuang, D.S., and Liao, J.C. (2021). Role of cyanobacterial phosphoketolase in energy regulation and glucose secretion under dark anaerobic and osmotic stress conditions. Metab Eng 65, 255–262.

De Porcellinis, A., Frigaard, N.-U., and Sakuragi, Y. (2017). Determination of the glycogen content in cyanobacteria. JoVE, e56068.

Diamond, S., Rubin, B.E., Shultzaberger, R.K., Chen, Y., Barber, C.D., and Golden, S.S. (2017). Redox crisis underlies conditional light–dark lethality in cyanobacterial mutants that lack the circadian regulator, RpaA. Proc Natl Acad Sci USA 114, E580–E589.

Evans, S.E., Franks, A.E., Bergman, M.E., Sethna, N.S., Currie, M.A., and Phillips, M.A. (2024). Plastid ancestors lacked a complete Entner-Doudoroff pathway, limiting plants to glycolysis and the pentose phosphate pathway. Nat Commun 15, 1102.

Field, C.B., Behrenfeld, M.J., Randerson, J.T., and Falkowski, P. (1998). Primary production of the biosphere: integrating terrestrial and oceanic components. Science 281, 237–240.

Flombaum, P., Gallegos, J.L., Gordillo, R.A., Rincón, J., Zabala, L.L., Jiao, N., Karl, D.M., Li, W.K.W., Lomas, M.W., Veneziano, D., Vera, C.S., Vrugt, J.A., and Martiny, A.C. (2013). Present and future global distributions of the marine Cyanobacteria *Prochlorococcus* and *Synechococcus*. Proc Natl Acad Sci USA 110, 9824–9829.

Hanson, A.D., Millar, A.H., Nikoloski, Z., and Way, D.A. (2023). Focus on respiration. Plant Physiol 191, 2067–2069.

Katayama, N., Iwazumi, K., Suzuki, H., Osanai, T., and Ito, S. (2022). Malic enzyme, not malate dehydrogenase, mainly oxidizes malate that originates from the tricarboxylic acid cycle in cyanobacteria. mBio 13, e02187–02122.

Kumar, S., Stecher, G., and Tamura, K. (2016). MEGA7: molecular evolutionary genetics analysis version 7.0 for bigger datasets. Mol Biol Evol 33, 1870–1874.

Lea-Smith, D.J., Bombelli, P., Vasudevan, R., and Howe, C.J. (2016). Photosynthetic, respiratory and extracellular electron transport pathways in cyanobacteria. Biochim. Biophys. Acta Bioenerg. 1857, 247–255.

Lu, K.-J., Chang, C.-W., Wang, C.-H., Chen, F.Y.H., Huang, I.Y., Huang, P.-H., Yang, C.-H., Wu, H.-Y., Wu, W.-J., Hsu, K.-C., Ho, M.-C., Tsai, M.-D., and Liao, J.C. (2023). An ATP-sensitive phosphoketolase regulates carbon fixation in cyanobacteria. Nat Metab 5, 1111–1126.

Mavridis, I.M., and Tulinsky, A. (1976). The folding and quaternary structure of trimeric 2-keto-3-deoxy-6-phosphogluconic aldolase at 3.5-A resolution. Biochemistry 15, 4410–4417.

Munson-McGee, J.H., Lindsay, M.R., Sintes, E., Brown, J.M., D’Angelo, T., Brown, J., Lubelczyk, L.C., Tomko, P., Emerson, D., Orcutt, B.N., Poulton, N.J., Herndl, G.J., and Stepanauskas, R. (2022). Decoupling of respiration rates and abundance in marine prokaryoplankton. Nature 612, 764–770.

Ohbayashi, R., Watanabe, S., Kanesaki, Y., Narikawa, R., Chibazakura, T., Ikeuchi, M., and Yoshikawa, H. (2013). DNA replication depends on photosynthetic electron transport in cyanobacteria. FEMS Microbiol Lett 344, 138–144.

Rust, M.J., Golden, S.S., and O’Shea, E.K. (2011). Light-driven changes in energy metabolism directly entrain the cyanobacterial circadian oscillator. Science 331, 220–223.

Shinde, S., Zhang, X.H., Singapuri, S.P., Kalra, I., Liu, X.H., Morgan-Kiss, R.M., and Wang, X. (2020). Glycogen metabolism supports photosynthesis start through the oxidative pentose phosphate pathway in cyanobacteria. Plant Physiol 182, 507–517.

Smith, A. (1983). Modes of cyanobacterial carbon metabolism. Ann. Inst. Pasteur Microbiol. 134, 93–113.

Tamoi, M., Miyazaki, T., Fukamizo, T., and Shigeoka, S. (2005). The Calvin cycle in cyanobacteria is regulated by CP12 via the NAD(H)/NADP(H) ratio under light/dark conditions. Plant J 42, 504–513.

Tanaka, K., Shirai, T., Vavricka, C.J., Matsuda, M., Kondo, A., and Hasunuma, T. (2022). Dark accumulation of downstream glycolytic intermediates initiates robust photosynthesis in cyanobacteria. Plant Physiol 191, 2400–2413.

Vizcaíno, J.A., Csordas, A., Del-Toro, N., Dianes, J.A., Griss, J., Lavidas, I., Mayer, G., Perez-Riverol, Y., Reisinger, F., and Ternent, T. (2015). 2016 update of the PRIDE database and its related tools. Nucleic Acids Res 44, D447–D456.

Wang, B., and Young, J.D. (2022). ^13^C-isotope-assisted assessment of metabolic quenching during sample collection from suspension cell cultures. Anal Chem 94, 7787–7794.

Watanabe, S., Ohbayashi, R., Shiwa, Y., Noda, A., Kanesaki, Y., Chibazakura, T., and Yoshikawa, H. (2012). Light-dependent and asynchronous replication of cyanobacterial multi-copy chromosomes. Mol Microbiol 83, 856–865.

Welkie, D.G., Rubin, B.E., Diamond, S., Hood, R.D., Savage, D.F., and Golden, S.S. (2018a). A hard day’s night: Cyanobacteria in diel cycles. Trends Microbiol.

Welkie, D.G., Rubin, B.E., Chang, Y.-G., Diamond, S., Rifkin, S.A., LiWang, A., and Golden, S.S. (2018b). Genome-wide fitness assessment during diurnal growth reveals an expanded role of the cyanobacterial circadian clock protein KaiA. Proc Natl Acad Sci USA 115, E7174–E7183.

Xiong, W., Lee, T.-C., Rommelfanger, S., Gjersing, E., Cano, M., Maness, P.-C., Ghirardi, M., and Yu, J. (2015). Phosphoketolase pathway contributes to carbon metabolism in cyanobacteria. Nat Plants 2, 15187.

Yu, H., Li, X., Duchoud, F., Chuang, D.S., and Liao, J.C. (2018). Augmenting the Calvin–Benson–Bassham cycle by a synthetic malyl-CoA-glycerate carbon fixation pathway. Nat Commun 9, 2008.

Zhang, S.Y., and Bryant, D.A. (2011). The tricarboxylic acid cycle in cyanobacteria. Science 334, 1551–1553.

Zhu, X.-G., Long, S.P., and Ort, D.R. (2008). What is the maximum efficiency with which photosynthesis can convert solar energy into biomass? Curr Opin Biotechnol 19, 153–159.

