## Supplementary information for "Phosphoketolase and KDPG aldolase metabolism modulate photosynthetic carbon yield in cyanobacteria"

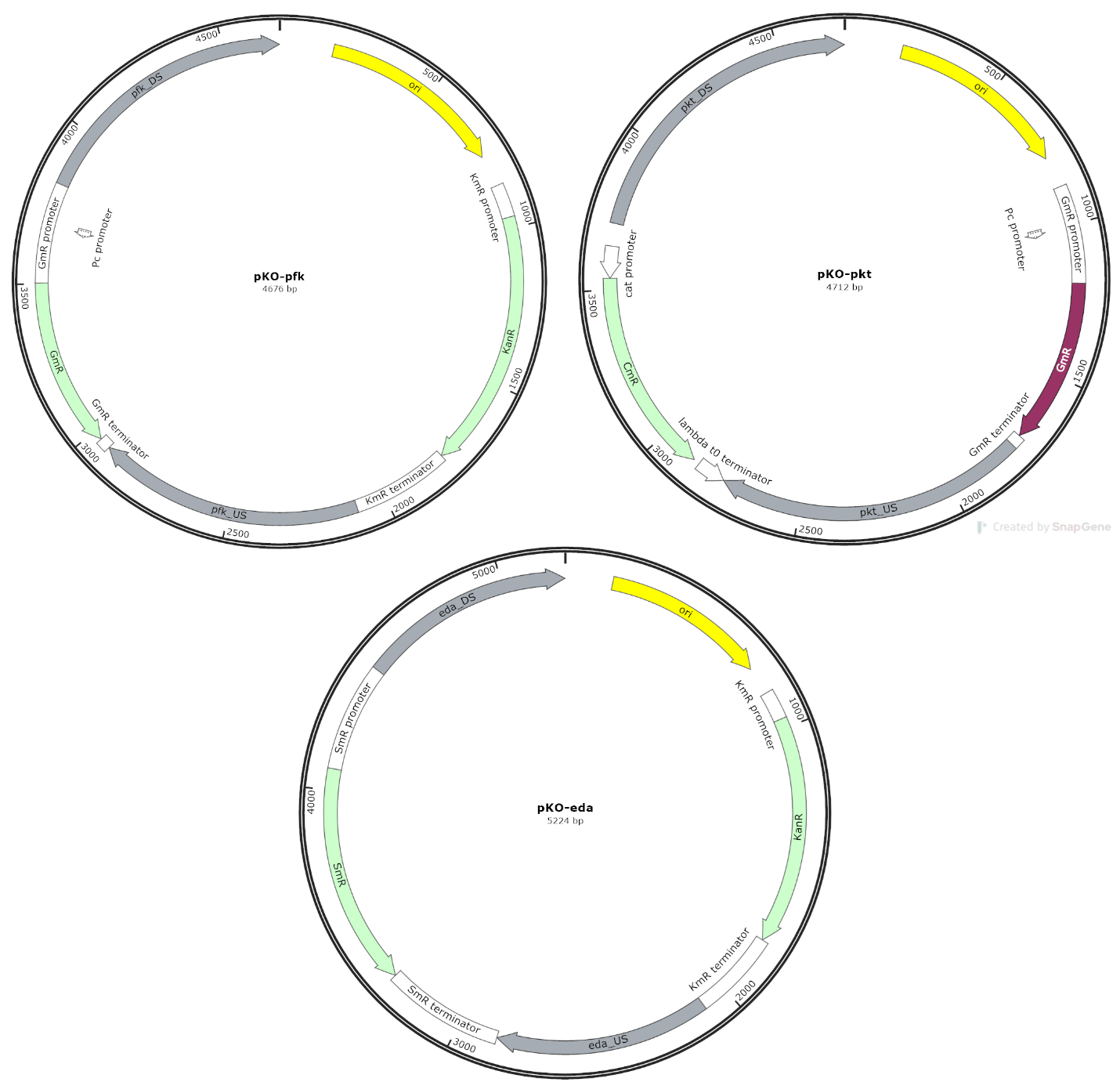

**Figure S1.** Plasmid maps for gene knockout. Phosphofructokinase (*pfk*), phosphoketolase (*pkt*), and ED aldolase (*eda*) were deleted from the genome of *Synechococcus elongatus* PCC 7942 wild type to construct mutant strains. This figure supports Figures 2, 3, 5, and 6 in the main manuscript.

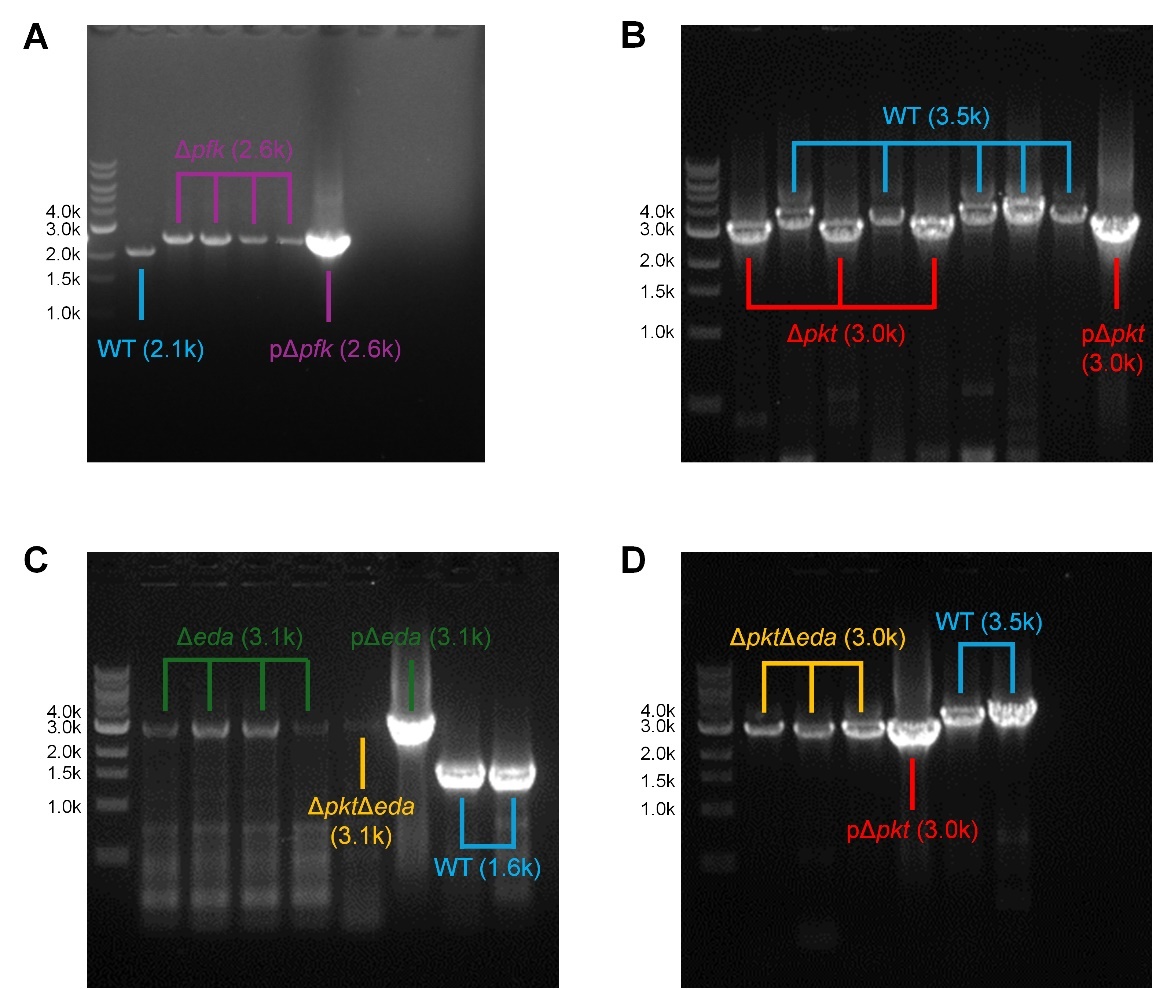

**Figure S2.** Segregation PCR for the mutant strains Δ*pfk*, Δ*pkt*, Δ*eda*, and Δ*pkt*Δ*eda*. **A)** The *pfk* gene is fully knocked out in the Δ*pfk* mutant. **B)** The *pkt* gene is fully knocked out in the Δ*pkt* mutant. **C)** The *eda* gene is fully knocked out in Δ*eda* and Δ*pkt*Δ*eda* mutants. **D)** The *pkt* gene is fully knocked out in the Δ*pkt*Δ*eda* mutant. Plasmids for constructing these mutant strains were used as positive controls (pΔ*pfk*, pΔ*pkt*, and pΔ*eda*) while the wild-type strain was used as the negative control (WT). This figure supports Figures 2, 3, 5, and 6 in the main text of the manuscript.

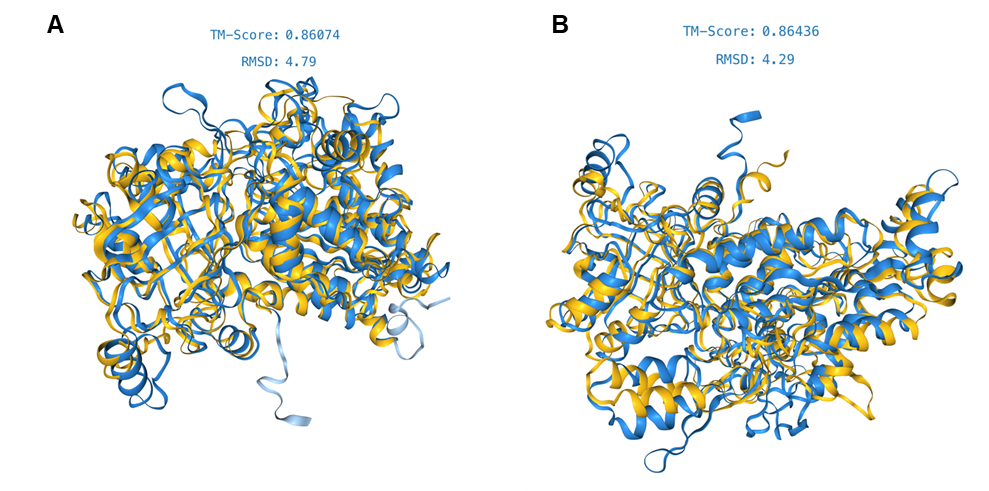

**Figure S3.** Fold Seek overlay *S. elongatus* DHAD with genuine 6PGDH protein from **(A)** *Escherichia coli* K10 and **(B)** *Pseudomonas aeruginosa*. To analyze and compare the structure of DHAD protein with 6PGDH from other organisms, Foldseek^1^ structure alignment was used. The TM score values were obtained from the analysis with score 1 showing perfect match and 0 to be a no match. This figure supports Figure 4 in the main manuscript.

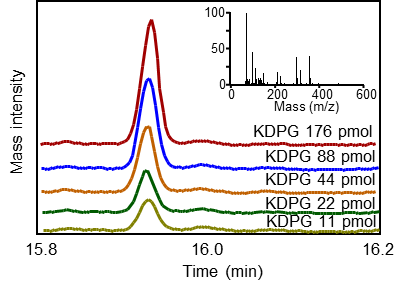

**Figure S4**. Detection of KDPG standards (ranging from 11 to 176 pmol) using GC-MS. The main plot shows the retention time of the KDPG standards. The inserted plot shows the mass spectrum of KDPG. This figure supports Figure 4 in the main text of the manuscript.

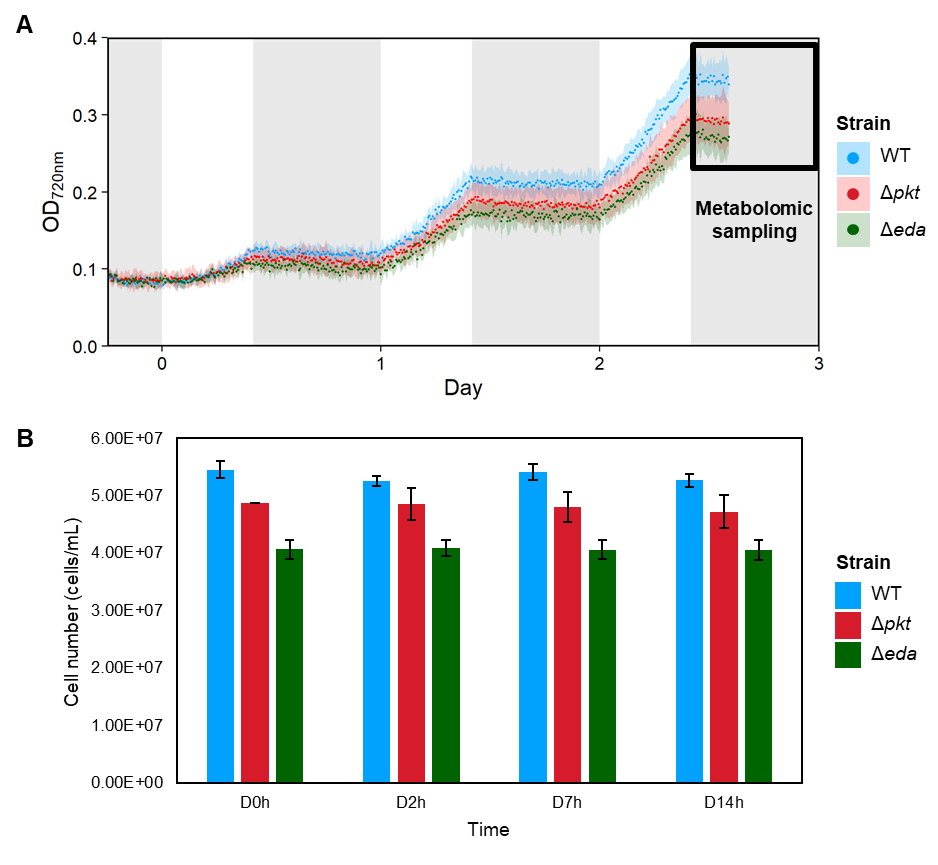

**Figure S5. Δ*pkt* and Δ*eda* show growth defects compared to the wild type during the diel cycles performed for metabolomic sampling.** **A)** The diel growth curve of each strain. The dotted lines and shades represent the OD_720nm_ mean and standard deviation of independently cultivated biological replicates (n = 5), respectively. The grey shading areas represent the dark periods. **B)** Cell numbers detected by flow cytometry at different time points in the dark period for metabolomic sampling. The error bar resents standard deviation of independently cultivated biological replicates (n = 5). This figure supports Figure 6 in the main text of the manuscript.

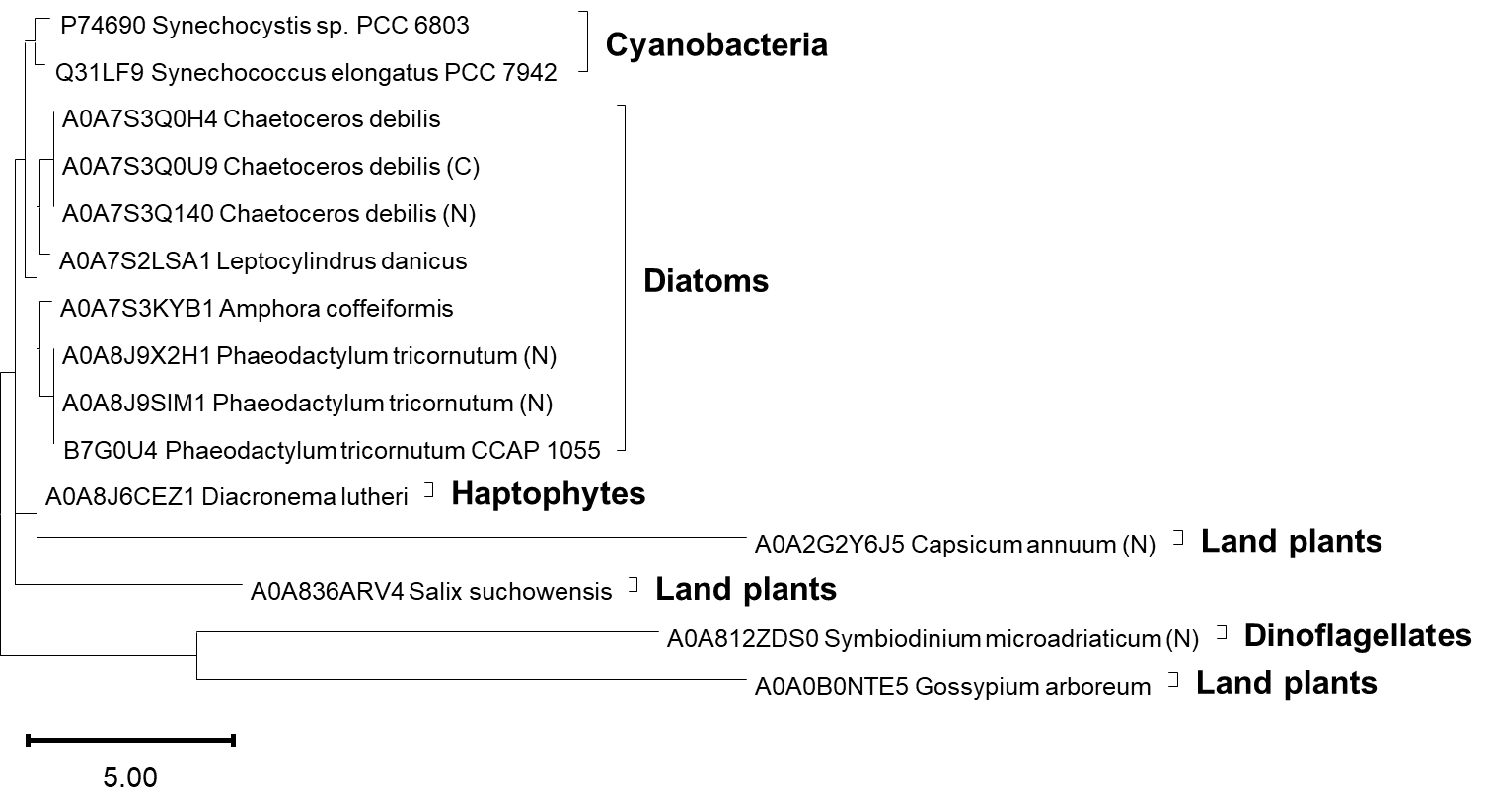

**Figure S6. Phylogenetic analysis of phosphoketolase in diverse photoautotrophic organisms.** Phosphoketolase exists across a broad range of photoautotrophic taxa including cyanobacteria, eukaryotic algae, and plants. The phylogenetic analysis was performed using protein sequences. C: C terminal; N: N terminal; otherwise, the whole phosphoketolase sequence was used. Phosphoketolase has been known to be prevalent in cyanobacteria and thus only two cyanobacterial strains were included as a reference. This figure supports Figure 7 in the main text of the manuscript.

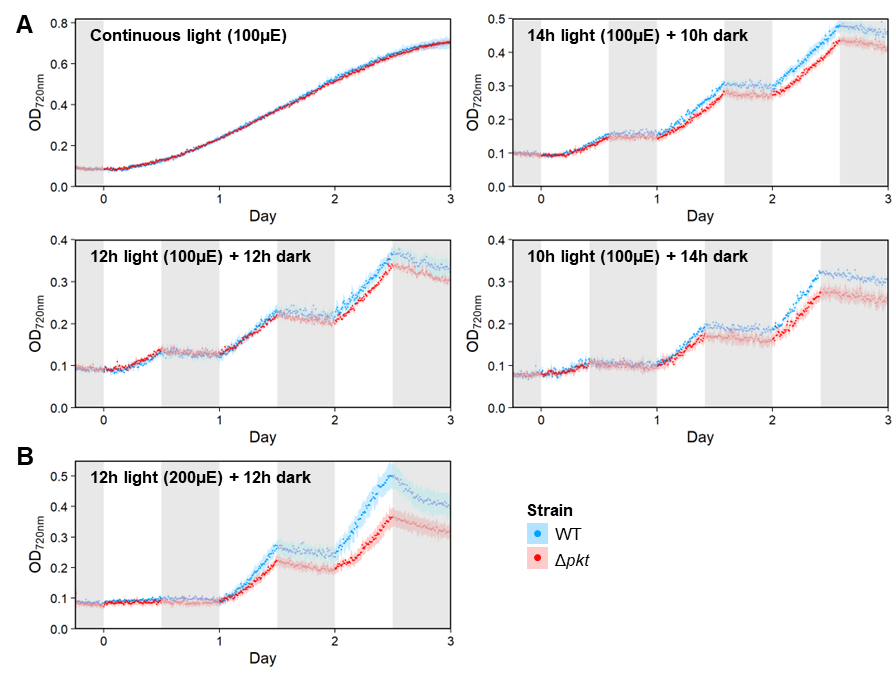

**Figure S7. Cyanobacterial growth under various light conditions.** Longer dark period and higher light intensity can exaggerate the diel growth defect of Δ*pkt* compared to the wild type. **A)** Growth of Δ*pkt* and WT in the continuous light and in the 10h/14h, 12h/12h, and 14h/12h light/dark cycles, using 100 µmol/m^2^/s of light for the light periods. **B)** Growth of Δ*pkt* and WT in the 12h/12h light/dark cycles, using 200 µmol/m^2^/s of light for the light period. This figure supports Figure 2 in the main text of the manuscript.

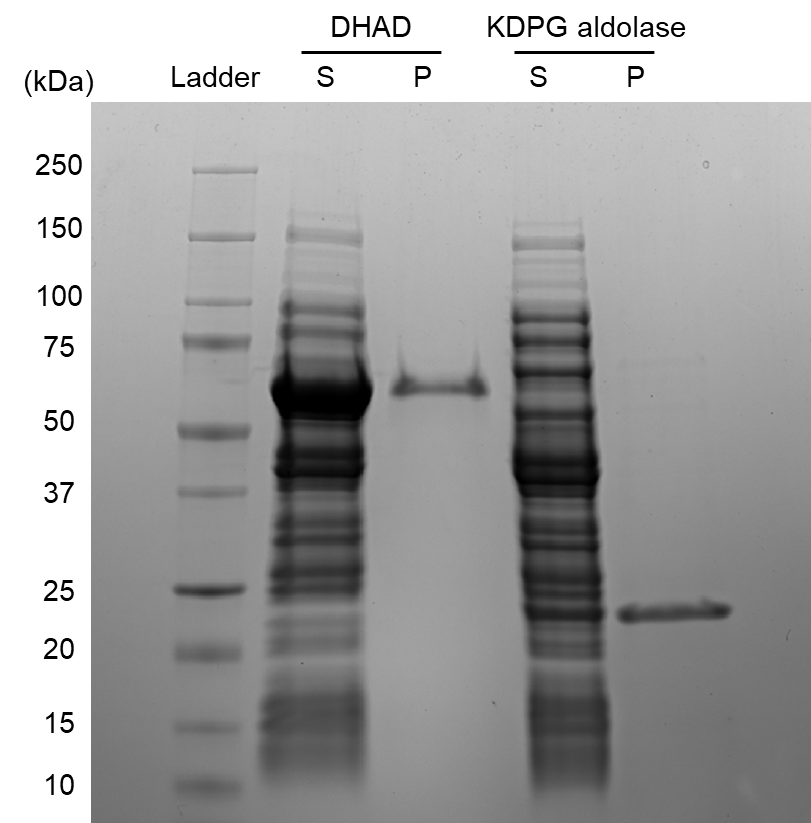

**Figure S8.** SDS-PAGE of recombinant DHAD and KDPG aldolase. S: supernatant; P: purified protein. This figure supports Figures 4 and 5 in the main text of the manuscript.

**Table S1.** Primers for plasmid and mutant strain construction. This table supports Figures 2, 3, 5, and 6 in the main text of the manuscript.

| Label | Sequence | Usage |
| --- | --- | --- |
| pKOpfk_Backbone_fwd | tccaaggccgatctgtcagaccaagtttactcatatatactttagattg | Knocking out *pfk* gene |
| pKOpfk_Backbone_rev | tgccgagcttggagatcgacctgcaggggg |  |
| pKOpfk_US_fwd | tgcaggtcgatctccaagctcggcaaggtcaatg |  |
| pKOpfk_US_rev | ttcccggccgcggcatcgccgccaatgccaatc |  |
| pKOpfk_GmR_fwd | ttggcggcgatgccgcggccgggaagccga |  |
| pKOpfk_GmR_rev | tgggcggtgattcgtggagaccgaaaccttgcgctcg |  |
| pKOpfk_DS_fwd | ttcggtctccacgaatcaccgcccaggatcc |  |
| pKOpfk_DS_rev | cttggtctgacagatcggccttggattgtgtttac |  |
| backbone-pkt_fwd | cggatgccagtgagcattcctcgcgcgactgtcagacc | Knocking out *pkt* gene |
| backbone-pkt_rev | tgtttggcctagcccgcggccgggaagccga |  |
| pkt_US_fwd | tcccggccgcgggctaggccaaacaactgc |  |
| pkt_US_rev | tattggtgagaatgtaagttccctcgagccaag |  |
| CmR_RC_fwd | gagggaacttacattctcaccaataaaaaacgc |  |
| CmR_RC_rev | gtcattgcttgcgagcgcaacgcaattaatg |  |
| pkt_DS_fwd | ttgcgttgcgctcgcaagcaatgaccaaggagc |  |
| pkt_DS_rev | cgcgaggaatgctcactggcatccgccctttag |  |
| Backbone_pKOeda_fwd | tccagagaatttctgtcagaccaagtttactcatatatactttagattg | Knocking out *eda* gene |
| Backbone_pKOeda_rev | caatgccctgctagatcgacctgcaggggg |  |
| eda_US_fwd | tgcaggtcgatctagcagggcattgattacg |  |
| eda_US_rev | acatatttgaatgcttcttccaaatccgtcag |  |
| SmR_pKOeda_fwd | atttggaagaagcattcaaatatgtatccgctcatgg |  |
| SmR_pKOeda_rev | ttctgtcggtgttgcgtggagaccgaaacc |  |
| eda_DS_fwd | cggtctccacgcaacaccgacagaaattatg |  |
| eda_DS_rev | cttggtctgacagaaattctctggattgggc |  |

**Table S2.** Statistical test results (*p*-values) on differences of metabolomic profiles between strains and time points. Pairwise comparisons were performed by permutation MANOVAs on the Bray-Curtis distance matrix after confirming equivalent beta-dispersion. The *p*-values were adjusted by FDR. This table supports Figure 6 in the main text of the manuscript.

A) ***p* values between strains**

|  | WT | Δ*pkt* |
| --- | --- | --- |
| Δ*pkt* | 0.001 |  |
| Δ*eda* | 0.001 | 0.001 |

B) ***p* values between time points**

|  | 0h | 2h | 7h |
| --- | --- | --- | --- |
| 2h | 0.162 |  |  |
| 7h | 0.006 | 0.009 |  |
| 14h | 0.006 | 0.006 | 0.494 |

**Table S3.** Coefficients of top 10 metabolites associated with each strain in the dark period (2h, 7h, and 14h), as revealed by Partial least squares discriminant analysis (PLS-DA). This table supports Figure 6 in the main text of the manuscript.

A) **Top 10 metabolites associated with the wild type.**

|  | WT | Δ*pkt* | Δ*eda* |
| --- | --- | --- | --- |
| chenodeoxycholic acid | 0.073 | -0.042 | -0.032 |
| 1-monoolein | 0.067 | -0.034 | -0.034 |
| 2-phosphoglyceric acid | 0.063 | -0.016 | -0.048 |
| pregnenolone | 0.058 | -0.035 | -0.024 |
| N-acetylglutamate | 0.057 | -0.115 | 0.057 |
| leucine | 0.050 | -0.024 | -0.027 |
| beta-hydroxymyristic.acid | 0.049 | 0.016 | -0.066 |
| 3-phosphoglycerate | 0.041 | -0.054 | 0.013 |
| maltotriose | 0.040 | -0.009 | -0.032 |
| quinic acid | 0.039 | -0.031 | -0.008 |

B) **Top 10 metabolites associated with Δ*pkt*.**

|  | WT | Δ*pkt* | Δ*eda* |
| --- | --- | --- | --- |
| adenine | 0.014 | 0.098 | -0.112 |
| 5’-deoxy-5’-methylthioadenosine | -0.040 | 0.084 | -0.043 |
| tyrosine | -0.031 | 0.081 | -0.050 |
| aspartate | -0.049 | 0.079 | -0.029 |
| threonine | -0.037 | 0.069 | -0.031 |
| lysine | -0.031 | 0.063 | -0.032 |
| isomaltose | -0.004 | 0.058 | -0.054 |
| ethanolamine | -0.104 | 0.057 | 0.049 |
| erythronic acid | -0.014 | 0.052 | -0.039 |
| oleic acid | -0.091 | 0.052 | 0.040 |

C) **Top 10 metabolites associated with Δ*eda*.**

|  | WT | Δ*pkt* | Δ*eda* |
| --- | --- | --- | --- |
| sarcosine | -0.061 | -0.032 | 0.094 |
| ribose-5-phosphate | -0.074 | -0.015 | 0.090 |
| oxoproline | -0.013 | -0.061 | 0.075 |
| trans-4-hydroxyproline | -0.009 | -0.063 | 0.072 |
| monolaurin | -0.033 | -0.036 | 0.069 |
| phthalic acid | -0.050 | -0.013 | 0.065 |
| glutathione | 0.022 | -0.086 | 0.063 |
| threose | -0.027 | -0.032 | 0.059 |
| N-acetylglutamate | 0.057 | -0.115 | 0.057 |
| 3-hydroxypalmitic acid | 0.009 | -0.065 | 0.056 |

**References**

1	van Kempen, M. *et al.* Fast and accurate protein structure search with Foldseek. *Nat. Biotechnol.*, doi:10.1038/s41587-023-01773-0 (2023).
